# The RPA complex orchestrates K63-linked deubiquitination via ZUP1

**DOI:** 10.1101/2023.01.27.525918

**Authors:** Benjamin Foster, Martin Attwood, Kay Chong, Matthew Drake, Ryan Gilroy, Marjorie Fournier, Ian Gibbs-Seymour

## Abstract

Ubiquitin signaling is regulated by deubiquitinating enzymes (DUBs), which can edit or remove ubiquitin from substrates. Recently, ZUP1 was discovered as the founding and only member of a novel DUB class that cleaves long K63-linked ubiquitin chains, with a putative role in DNA repair. However, ZUP1 has poor activity on its own, suggesting additional mechanisms exist to promote its activity in cells. Here, using a range of cellular, biochemical, and structural proteomics approaches, we show that ZUP1 directly interacts with the RPA complex, a single-stranded DNA binding protein complex involved in DNA replication and multiple DNA repair processes. Functionally, the ZUP1-RPA complex interaction dramatically stimulates ZUP1 K63-linked DUB activity, which occurs through S1 and S1’ site communication. Collectively, our results suggest a mechanism that couples sensing of ssDNA to the activation of K63-linked deubiquitination via the ZUP1-RPA complex.

## Introduction

The covalent attachment of ubiquitin (Ub) to substrates is a post-translational modification (PTM) that has a wide-ranging role in regulating myriad cellular pathways, including proteostasis, inflammation, and the DNA damage response (DDR) ^1–3^. The addition of Ub to substrates occurs via a number of enzymatic steps involving E1-activating, E2-conjugating, and E3-ligating enzymes. Canonical Ub attachment to substrates is via an isopeptide bond linking the C-terminal carboxyl group of Ub with the *ε*-amino group of a lysine residue in the target protein substrate ^4^. Additionally, the *α*-amino group of Ub Met1 (M1) can also act as a donor for Ub attachment ^5^. There is also growing evidence for non-lysine Ub attachment modes, through oxyester (to serine and threonine) or thioester (to cysteine) bond formation ^6, 7^. Furthermore, non-proteinaceous substrate targets have also recently emerged, such as ADP-ribose, lipids, and sugars ^8–11^. Thus, the different modes of attachment and scope of possible macromolecular substrates has recently expanded the potential for Ub signaling in cells. Moreover, as Ub contains seven internal lysine residues (K6, K11, K27, K29, K33, K48, and K63), together with the M1 residue, it can serve as attachment sites for another Ub molecule, enabling the formation of polymeric Ub chains (polyUb) ^12^. Importantly, polyUb of different linkage types can be decoded by effector proteins with Ub binding domains (UBDs), which translate the Ub code into functional outcomes and are associated with different signaling pathways ^12–14^. Taken together, the expanded scope of possible substrates, the novel modes of Ub attachment, and the ability to generate complex polyUb architectures, highlights the vastness of the Ub code. Yet set against this, Ub signaling can be highly specific and dynamic to direct cell fate outcomes.

For ubiquitination to be highly dynamic in cellular signaling pathways, it should be reversible. The maturation of Ub precursor proteins and the removal of Ub and polyUb chains from substrates is carried out by dedicated proteases called deubiquitinases (DUBs) ^15, 16^. There are approximately 100 DUBs encoded in the human genome that can be classified into seven distinct families based on their structural features and evolutionary ancestry ^17^. DUB activity depends on their ability to recognise the distal Ub via their S1 site, which presents the C-terminus of Ub to the active site for hydrolysis, providing specificity for Ub versus other Ub-like modifiers ^15, 18^. In addition, many DUBs also have an additional binding site that interacts with the proximal Ub, termed the S1’ site, which helps define the linkage specificity. Further Ub-binding sites can also contribute to Ub recognition, and are termed S2, S3, S2’, S3’ sites. As important regulators of the Ub system, DUBs are becoming increasingly implicated in human disease and, consequently, there has been much recent interest in exploring DUBs as potential therapeutic targets ^18, 19^. Thus, further understanding the biology and mechanisms of DUBs will be essential moving forward.

Zinc finger-containing Ub peptidase 1 (ZUP1) was recently discovered as the founding, and so far only member, of a seventh DUB class that cleaves long K63-linked polyUb chains ^20–23^ (Extended Data Figure 1A). Whilst the catalytic residues reside in the C-terminal C78 peptidase domain, the domain in isolation is unable to cleave Ub chains ^20–23^. Rather, ZUP1 DUB activity and specificity for longer K63-linked polyUb chains is stimulated by several additional UBDs in the protein ^20–23^. Adjacent to the C78 domains is an atypical Ub binding motif termed the ZUP1 helical arm (ZHA), which binds the distal S1 Ub ^20, 22^ (Extended Data Figure 1B). A motif interacting with ubiquitin (MIU) domain next to the ZHA is the likely S2 binding site in a K63-linked polyUb chain. ZUP1 also contains four zinc finger domains (ZnFs), which are similar at the sequence level to Ub binding zinc finger (UBZ) domains. One of these, ZnF4 (which we refer to as the UBZ), is particularly important for overall polyUb binding in ZUP1 ^20–22^ (Extended Data Figure 1C). The UBZ might bind the S3 (or beyond) Ub in a K63-linked polyUb chain, distal to the C78 domain, or it may form part of the S1’ Ub binding site proximal to the C78 domain, depending on its relative orientation – these two models of polyUb chain recognition by ZUP1 await to be empirically determined (Extended Data Figure 1D,E). Lastly, a helix-turn-helix motif (annotated as *α*-2*/*3 in the crystal structures) was also found to be required for ZUP1 DUB activity, likely acting as part of the S1’ binding site involved in stabilising the proximal Ub across the active site ^20, 22, 24^. Since the published ZUP1 structures lacked the UBZs and contained only the monoubiquitin bound to the S1 site, the contributions of the multiple UBDs in ZUP1 to substrate recognition and how it achieves K63-linkage specificity remain unknown. Despite the presence of several UBDs, ZUP1 is a poor enzyme on its own *in vitro*, suggesting that there might be additional mechanisms to stimulate its activity within a cellular context ^20–23^.

**Figure 1.**
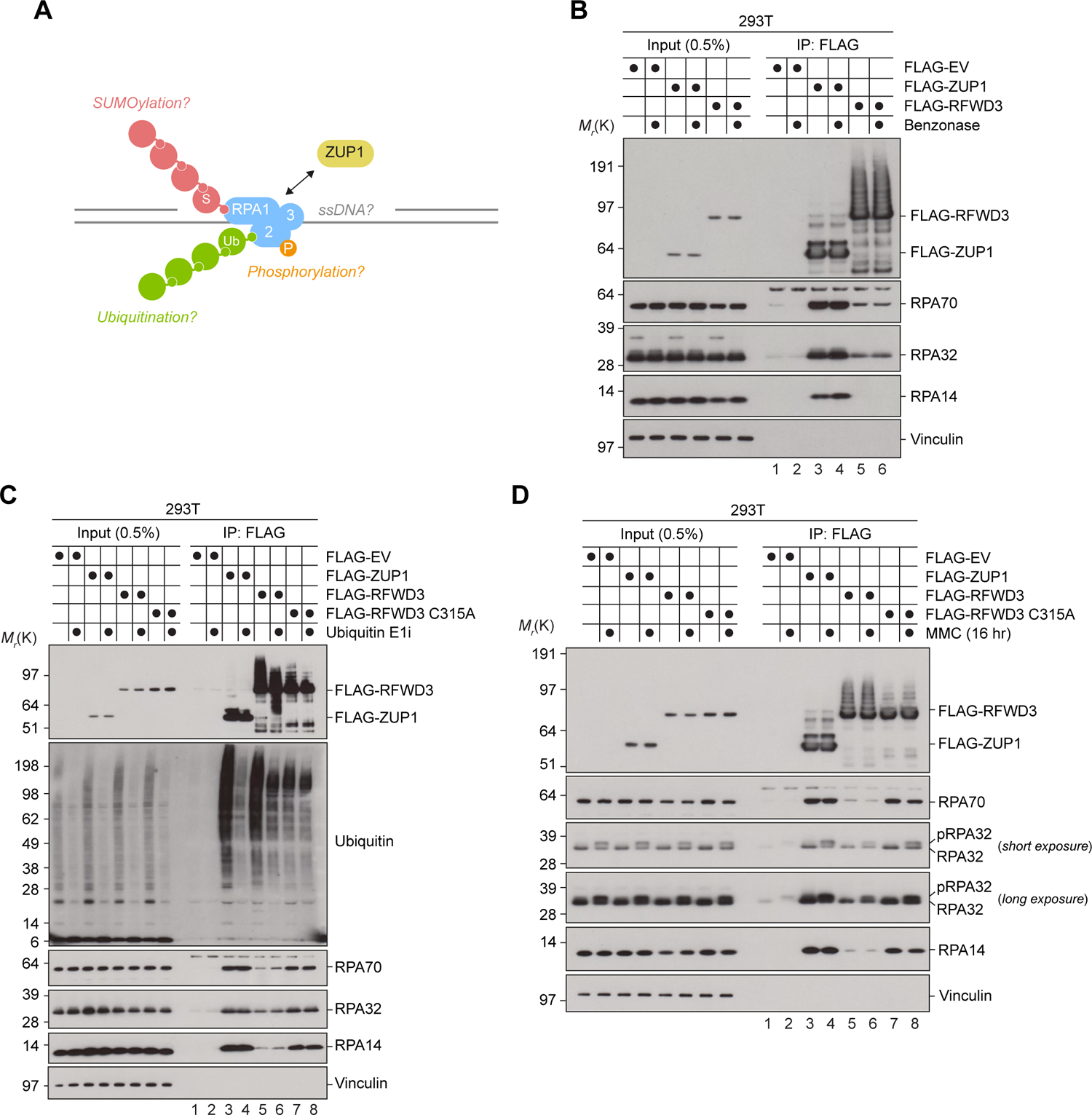
ZUP1 and RPA form a complex in human cells. **A.** Schematic showing the variables that were assessed during our characterisation of the ZUP1-RPA interaction in cells. **B.** 293T cells were transfected with FLAG-empty vector (EV), FLAG-ZUP1 or FLAG-RFWD3. Cells were lysed in the absence (-) or presence (+) on benzonase (>500 units/ml), subjected to FLAG immunoprecipitation (IP), and analysed by immunoblotting with the indicated antibodies. Data is representative from *n* > 3 biological repeats. **C.** 293T cells were transfected with FLAG-EV, FLAG-ZUP1, FLAG-RFWD3, or FLAG-RFWD3 C315A and then treated with 1 µM Ub E1i for 90 mins. Cells were lysed, subjected to FLAG immunoprecipitation (IP), and analysed by immunoblotting with the indicated antibodies. Data is representative from *n* = 2 biological repeats. **D.** As for (C), except cells were treated with 1 µM MMC for 16 h before lysis. Data is representative from *n* = 2 biological repeats.See also Extended Data Figure 1.

Ub signaling plays an important role in multiple DNA repair pathways, with some ubiquitination targets being critical signaling events for those pathways. Examples include the ubiquitination of the FANCD2-FANCI complex in the Fanconi Anaemia (FA) pathway, ubiquitination of PCNA in translesion synthesis (TLS), template switching, and fork reversal, and ubiquitination of replication protein A (RPA), which is involved in coordinating DNA replication and multiple DNA repair pathways ^3^. RPA is a heterotrimeric single-stranded DNA (ssDNA) binding complex, consisting of RPA70, RPA32, and RPA14 subunits ^25^. Within the RPA complex, there are multiple oligonucleotide/oligosaccharide (OB)-fold ssDNA binding domains distributed across the subunits, separated by flexible linker regions, which provide the complex with an ability to bind ssDNA with high affinity (K_d_, 10^-10^M) and with multiple possible conformations ^26–28^. Thus, RPA can bind ssDNA that is generated in the cell, preventing potentially deleterious secondary structure formation or nucleolytic processing of the ssDNA. An additional essential function of RPA is to coordinate the recruitment of numerous DNA replication and DNA repair factors via protein-protein interactions, which typically occurs through two main regions in RPA, the N-terminal domain of RPA70 (called OB-F or PID^70N^, for protein interaction domain) or the C-terminal winged helix domain of RPA32 (termed WH or PID^32C^) ^29, 30^. One prominent example of an RPA interaction is the ATR-ATRIP kinase complex that triggers checkpoint signaling in response to ssDNA generation. In addition, RPA also coordinates Ub signaling at ssDNA via interaction with a number of Ub E3 ligases, such as RAD18 ^31^, RFWD3 ^32–35^, PRP19 ^36^, and HERC2 ^37^. Whilst there is mounting evidence for RPA coordinating ubiquitination events, there is little evidence of how RPA might coordinate deubiquitination events at ssDNA, despite numerous DUBs being implicated in the DDR ^38^. Moreover, for most DUBs it is unclear how they achieve substrate specificity and how their activities are regulated. ZUP1 was previously implicated in the DDR and was shown to interact with several proteins involved in DNA replication and DNA repair, including RPA ^20–23, 39^. However, it was unclear whether this interaction was direct or mediated by DNA and/or other protein(s) and there was no further mechanistic detail beyond the observed cellular interaction. Moreover, it was unknown whether the interaction had any functional relevance to the biochemical activity of ZUP1.

Here we provide evidence of a mechanism by which a DUB is activated in a context-dependent manner. Specifically, we show that ZUP1 directly interacts with the RPA complex, both in cells and *in vitro*. We map the RPA binding site to the ZUP1 *α*-2*/*3 region, the proposed S1’ site in ZUP1. Importantly, we find that the interaction with RPA stimulates the ZUP1 DUB activity against K63-linked polyUb chains. Mechanistically, the binding of RPA to the *α*-2*/*3 region acts at least through the adjacent ZHA S1 site in ZUP1, suggesting that the RPA interaction promotes ZUP1-Ub formation. We further show that the stimulation of ZUP1 DUB activity by RPA is both specific to RPA and evolutionarily conserved. Overall, our findings suggest that the RPA complex orchestrates K63-linked deubiquitination events at ssDNA via its interaction with, and stimulation of, ZUP1.

## Results

### ZUP1 and RPA form a complex in human cells

We and other groups previously reported a cellular interaction between ZUP1 and RPA, which was discovered either by mass spectrometry and/or co-immunoprecipitation analyses from human cells ^20–23^. One study showed that the ability of FLAG-ZUP1 to co-immunoprecipitate RPA from human cells is lost after nuclease addition to the cell lysis buffer, which led them to suggest that the interaction might be mediated by DNA, either through RPA and/or ZUP1 DNA binding, and/or by another bridging protein(s) ^22^. Thus, the details of the ZUP1-RPA complex interaction remained largely opaque. We reasoned that defining the interaction between ZUP1 and RPA was important because it should provide further mechanistic details on this DUB class, such as how ZUP1 is recruited to DNA lesions, how its activity might be regulated, and how it chooses substrates to deubiquitinate. Given that RPA is the target of various PTMs (Figure 1A) ^30^, we therefore sought to systematically analyse the ZUP1-RPA interaction in cells under a range of conditions related to RPA function, such as DNA binding, RPA phosphorylation, ubiquitination, and SUMOylation. For these analyses, we used the Ub E3 ligase RFWD3 as a positive control, as it has been shown to directly interact with RPA ^40, 41^. Firstly, to re-examine the previous observation that the ZUP1-RPA interaction might be DNA-dependent, we expressed FLAG-ZUP1 or FLAG-RFWD3 in 293T cells and immunopurified each of them from cell lysates which were prepared in the presence or absence of the benzonase nuclease. In parallel, nucleic acids from the same lysates were analysed by agarose gel electrophoresis to ensure benzonase treatment was successful and genomic DNA was digested (Extended Data Figure 1F). Immunoblot analysis of the immunopurifications showed that FLAG-ZUP1 co-purified all three RPA subunits (RPA70, RPA32 and RPA14) in the presence or absence of benzonase, indicating that the ZUP1-RPA interaction does not depend on the presence of DNA (Figure 1B, compare lanes 3 and 4). Similarly, RFWD3 also co-purified RPA, albeit to a far lesser extent than ZUP1, which was also independent of benzonase treatment and therefore genomic DNA, which was expected given the previous evidence of a direct interaction between them (Figure 1B, compare lanes 5 and 6). Whilst we cannot exclude the possibility that smaller fragments of DNA retained after benzonase treatment might bridge the interaction between ZUP1 and RPA, the above data suggested that the ZUP1-RPA interaction is likely independent of DNA.

Next, as RPA has been reported to be ubiquitinated by RFWD3 and other E3 ligases, we examined whether inhibiting the Ub system using the TAK-243 Ub E1 inhibitor (E1i) impacted the ZUP1-RPA interaction. Cells treated with the Ub E1i showed dramatically reduced polyUb levels, as evidenced by the loss of Ub signal in the inputs and in the signal of the proteins co-purified with either ZUP1 or RFWD3 (Figure 1C, compare odd and even numbered lanes). However, inhibiting ubiquitination had no impact on the ability of FLAG-ZUP1 or FLAG-RFWD3 to co-purify the RPA complex (Figure 1C, compare lanes 3-6). Here, we also used the RFWD3 C315A mutant, which renders RFWD3 catalytically inactive as an E3 ligase and also acts as a substrate trapping mutant ^34^. The RFWD3 C315A mutant was able to co-purify far more RPA complex than RFWD3 wild type (WT), at levels more comparable to ZUP1 (Figure 1C, lanes 3, 4, 7, and 8). Notably, the Ub E1i also impaired the slower migrating species of FLAG-ZUP1 and FLAG-RFWD3 WT, suggesting that these species are polyubiquitinated forms of the two proteins (Figure 1C, compare lanes 4 and 6). For RFWD3 C315A, this FLAG signal is already compromised, indicative of impaired autoubiquitination by this mutant (Figure 1C, lane 8). We conclude from these experiments that the ZUP1-RPA interaction can largely occur independently of Ub signaling in cells. As RPA is also a target for the small ubiquitin-like modifiers (SUMO), we analysed the ZUP1-RPA interaction after treating cells with the SUMO E1i ML-792. Similar to the Ub E1i, we found that inhibition of SUMO signaling had negligible impact on the ability of ZUP1 to co-purify RPA from cells (Extended Data Figure 1G).

Lastly, we examined the ZUP1-RPA interaction after treatment of cells with the DNA damaging drug mitomycin C (MMC), a chemotherapeutic that causes interstrand crosslinks (ICLs) and activates the FA pathway, as it was previously shown that RFWD3-deficient cells have defects in ICL repair ^33, 34^. Under conditions of MMC treatment, RPA undergoes extensive phosphorylation mainly through the RPA2 subunit, resulting in slower migrating forms in SDS-PAGE analyses. We found that after DNA damage via MMC, ZUP1 could co-purify the phosphorylated RPA complex (Figure 1D) and that this phosphorylation mainly occurred through the ATR kinase (Extended Data Figure 1H). Taken together, our data suggest that ZUP1 and RPA form a constitutive complex in cells.

A limitation to these experiments is that both ZUP1 and RFWD3 are over-expressed in cells, which may potentially saturate endogenous regulation. However, the anti-ZUP1 antibodies we have tested have been suboptimal for co-immunoprecipitation experiments. Endogenous tagging of ZUP1 is also complicated by the number of isoforms expressed in cells. Despite this, we do not observe any clear differences in the ZUP1-RPA interaction under all the conditions tested here. Thus, the data from these cellular assays suggested a more parsimonious explanation for prior observations: that the ZUP1-RPA complex interaction might be direct and, if so, might impact the biochemical activity of ZUP1.

### ZUP1 and RPA form a complex in vitro

To determine if ZUP1 can directly bind to RPA, we prepared purified recombinant full-length ZUP1 and full-length RPA from *E. coli*. Firstly, we observed that recombinant RPA robustly co-purifies with a GST-tagged ZUP1 in GST-pulldown experiments when analysed by SDS-PAGE and protein staining with Coomassie (Figure 2A). To extend this finding beyond immobilised ZUP1, we used size exclusion chromatography (SEC) analysis to show that the ZUP1-RPA interaction also occurs in solution, in the absence of ssDNA, with a clear peak shift on the chromatogram, which was confirmed by analysis of the corresponding fractions via SDS-PAGE (Figure 2B). Our findings therefore extend and support our observations from cells, that ZUP1 and RPA form a complex via a direct interaction(s).

**Figure 2.**
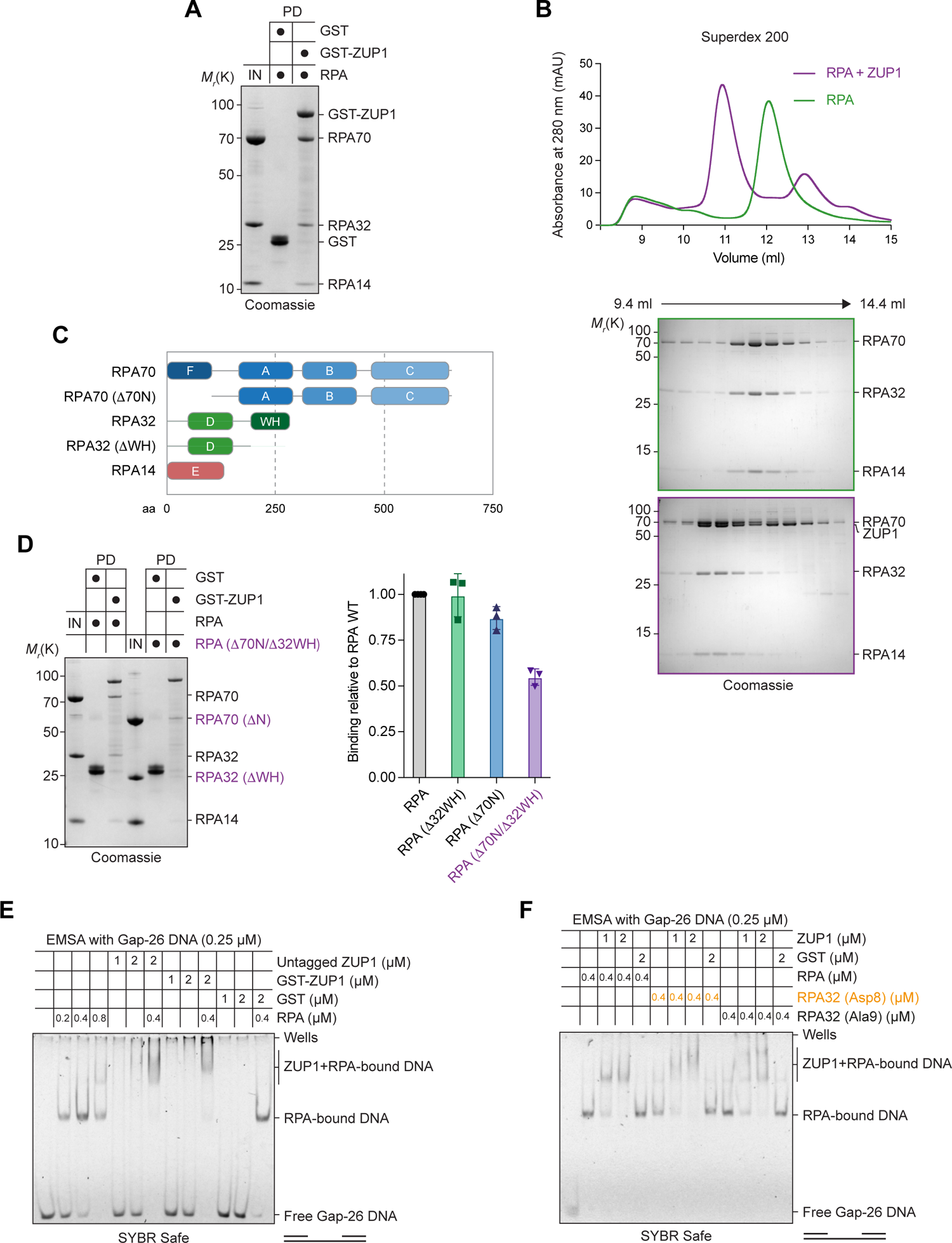
ZUP1 and RPA form a complex *in vitro*. **A.** Recombinant GST or GST-tagged ZUP1 were pre-bound to glutathione beads and then incubated with purified untagged RPA complex for 1 h. Bound material was eluted, separated by SDS-PAGE gel, and then stained with Coomassie Brilliant Blue. 20% of the RPA input (IN) and 50% of the eluted pulldown (PD) was used. **B.** Chromatogram from analytical SEC showing the absorbance at 280 nm of RPA and ZUP1 or RPA alone or run over a Superdex 200 10/300 gel filtration column. Proteins were eluted and peak fractions were separated by SDS-PAGE gel and stained with Coomassie Brilliant Blue. 300 µg of each were mixed and loaded, resulting in a ∼1.7 molar excess of ZUP1 over RPA. Free ZUP1 can be detected in the second, smaller peak fractions indicating approximately 1:1 stoichiometry. **C.** Domain diagram of the RPA complex, indicating the mutants used here. **D.** *Left*, as in (A), except with the indicated RPA complex deletion mutant. *Right*, quantification of the bands corresponding to RPA70 (WT, *Δ*32WH, *Δ70*N, or *Δ70*N/*Δ*32WH) in each pulldown. Data is from *n* = 3 independent experiments and represents the mean and standard deviation. **E.** EMSA using a Gap-26 unlabelled DNA substrate with the indicated molarities of recombinant ZUP1, GST, and RPA complex. The migration of DNA was analysed by 5% native PAGE and staining with SYBR Safe. **F.** As for (D), except phospho-mimetic (32Asp8) or phospho-null (32Ala9) RPA complex was used. See also Extended Data Figure 2.

We next sought to identify the region(s) in RPA responsible for the ZUP1 interaction. To define the RPA subunit(s) involved, we found that GST-ZUP1 could co-purify untagged recombinant RPA70 and RPA32 when individually expressed from *E. coli* cell lysates, but not RPA14 (Extended Data Figure 2A). RPA has two main protein-protein interaction domains, the RPA70 N-terminal OB-F domain and the RPA32 C-terminal WH domain, with the majority of RPA interacting factors binding via one or both of these domains (Figure 2C) ^42^. We therefore produced mutant RPA by deleting the RPA70 N-terminal OB-F domain (*Δ*70N) and RPA32 C-terminal WH domain (*Δ*32WH), either singly or together, and analysed the interaction with ZUP1. Whilst deletion of each individual domain had negligible impact, the double deletion (*Δ*70N/*Δ*32WH) led to a ∼50% reduction in the ability of GST-ZUP1 to co-purify RPA (Figure 2C and Extended Data Figure 2B). Notably, the isolated RPA70 N-terminal region, but not the RPA32 C-terminal WH, could co-purify ZUP1 in GST pulldown assays (Extended Data Figure 2C). From this data, we conclude that ZUP1-RPA complex formation is likely promoted through multiple interactions in RPA and is only partially dependent on the RPA70 OB-F and RPA32 WH domains.

We next examined ZUP1-RPA complex formation in the context of RPA ssDNA binding and RPA phosphorylation, given that these are critical to RPA function. Electrophoretic mobility shift assay (EMSA) using a ssDNA-containing DNA substrate suggested that the ZUP1-RPA complex can bind ssDNA (Figure 2E). We examined numerous other DNA substrates with ssDNA regions that confirmed this observation (data not shown). Moreover, we found that the interaction is maintained in the presence of dt100 ssDNA using SEC analysis (Extended Data Figure 2D). Using an RPA mutant with Ser to Asp mutations in the N-terminus of RPA32 (called Asp8), which mimics the RPA32 hyper-phosphorylation that occurs after ssDNA generation in cells, or an RPA mutant with Ser to Ala mutations (called Ala9), we observed no difference in the band shifts of either of these two ZUP1-RPA-DNA complexes compared to RPA WT, when analysed by EMSA (Figure 2F). This indicates that ZUP1 can directly bind to the RPA complex on ssDNA independently of RPA32 N-terminal hyper-phosphorylation, at least using these phospho-mimetic conditions. Collectively, our data suggest that the ZUP1 and RPA form a complex *in vitro* that binds ssDNA.

### The ZUP1-RPA complex contains multiple contact sites

The data so far argues for a ZUP1-RPA complex that can bind ssDNA, thereby linking ssDNA detection by RPA to deubiquitination by a Ub K63-specific DUB. Thus, understanding the structural basis for this interaction is a key aim. However, despite our extensive efforts, we are yet to obtain a cryo-EM structure of the ZUP1-RPA complex either with or without ssDNA and/or in combination with a K63-linked polyUb chain. Obtaining crystal structures for the full-length RPA complex has been historically difficult, given the flexible linkers between the various domains. Indeed, the most detailed crystal structure of the RPA complex to date was obtained by removing the flexible linkers between ^43^. A more recent cryo-EM structure of the RPA complex on ssDNA was also missing substantial parts of the complex ^44^. In addition, ZUP1 has a long flexible linker between the ZnF2 and ZnF3 domains, providing further difficulty for structural approaches with full-length ZUP1. To gain further insight into the full-length ZUP1-RPA complex, we developed a crosslinking with mass spectrometry (XL-MS) workflow (Extended Data Figure 3A). The ZUP1-RPA complex was formed *in vitro*, then treated with the lysine-reactive crosslinker bis(sulphosuccinimidyl)suberate (BS3), with subsequent processing and analysis by mass spectrometry. Alternatively, samples were subjected to a strong cation exchange (SCX) chromatography step to enrich for highly charged crosslinked peptides prior to analysis by mass spectrometry (Extended Data Figure 3B and Supplementary Data 1 and 2). As expected, the SCX step improved the ability to detect crosslinks within the ZUP1-RPA complex, especially between ZUP1 and RPA14, though in general the crosslinked regions were largely similar (Figure 3A,B). Importantly, we detected multiple crosslinks between ZUP1 and regions on both RPA70 and RPA32 (Figure 3A,B). The crosslinks between ZUP1 and RPA32 were focussed on the C-terminal WH domain, whilst there was a more even distribution of crosslinks between ZUP1 and RPA70 across both proteins. We also detected crosslinks between ZUP1 and RPA via regions proximal to the C78 peptidase domain and the *α*-2/3 region, specifically on K262 and K522/525 in ZUP1 (see Figure 3C for residue locations). Notably, the disordered region between the ZnF2 and ZnF3 domains in ZUP1 was a hotspot for crosslinks, likely due to its flexibility and the density of lysine residues located there. Together, the XL-MS data show that extensive multi-site interactions between ZUP1 and RPA stabilise the complex. This is highlighted by the RPA *Δ*70N/*Δ*32WH mutant which lacks the two canonical protein-protein interacting domains in RPA, which still shows binding to ZUP1in pull downs.

**Figure 3.**
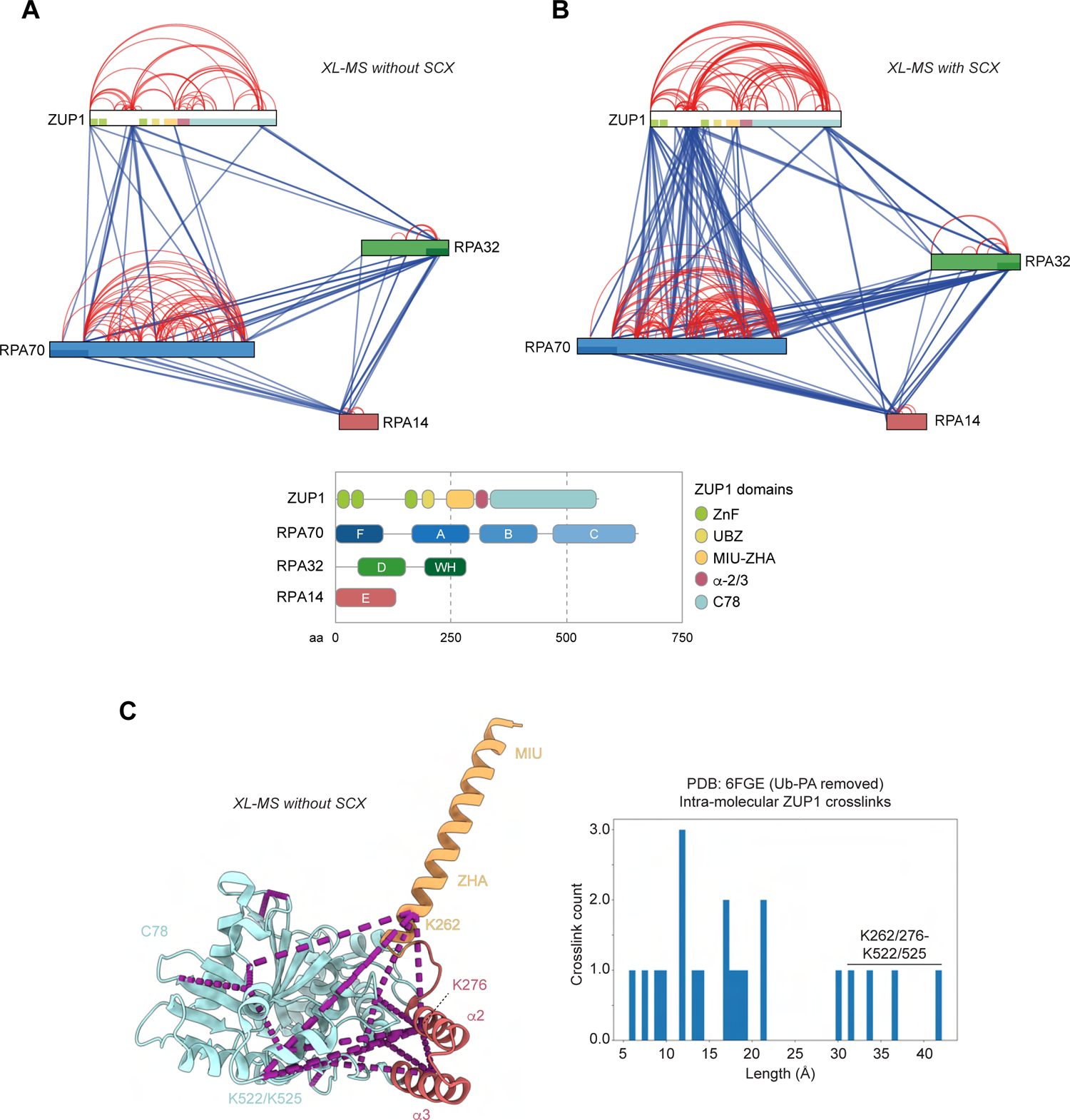
XL-MS analysis of the ZUP1-RPA complex. **A.** Recombinant ZUP1 and RPA were crosslinked with BS3, processed for mass spectrometry, and then analysed with pLink2.0, with identified crosslinks visualised using xVis. Domain colours within the polypeptides in the interaction network correspond to the domain colours in the domain diagrams below. Inter-molecular crosslinks are shown in blue and intra-molecular crosslinks in red. Only crosslinks detected in both replicates (>1 technical replicate) and/or with multiple spectral evidence present were included. **B.** As in (A), but with an additional strong cation exchange (SCX) step during the sample processing. **C.** *Left*, the structure of ZUP1 aa 231-578 (PDB: 6FGE) with intramolecular crosslinks indicated by purple dashed lines. *Right*, histogram showing distance measurements of the intramolecular crosslinks detected in ZUP1. BS3 is likely to enable crosslinks at a maximum of ∼30 Å (11 Å of the crosslinker, plus two lysine side chains), so the majority are within this distance. Those above this limit all occur between the ZHA and *α*-2/3 region to K522/525, which indicates conformational flexibility of this region. See also Extended Data Figure 3.

The XL-MS data also showed long-range interactions within ZUP1, with crosslinks from the ZnFs and the disordered region preceding ZnF3 to the C-terminus (Figure 3A,B). Moreover, we also observe MIU-ZHA crosslinks to the C-terminus. When these were mapped on to the ZUP1 crystal structure with Ub-PA bound to the active site Cys360 (PDB: 6FGE), the distances of the crosslinks between the ZHA (K262/K276) and the C-terminus (K522/K525) exceed the maximum distance expected for the BS3 crosslinker, suggesting some degree of conformational flexibility to allow formation of these crosslinks (Figure 3C). Whether these and the longer N-to-C-terminal crosslinks are truly intra-molecular long-range interactions cannot be definitively determined using this XL-MS approach. However, it is notable that ZUP1 forms a monomer in solution at the molarities used in these experiments when analysed by gel filtration and SEC-MALS (Extended Data Figure 3C,D), and the ZUP1-RPA complex has a 1:1 stoichiometry as assessed by analytical SEC (Extended Data Figure 3C), indicating the crosslinks occur between domains within a ZUP1 monomer rather than between different ZUP1 protomers. Collectively, the XL-MS data supports our prior biochemical findings presented above and extends them by indicating multiple contact points within the ZUP1-RPA complex, which may be relevant for the overall formation of the complex. Moreover, the XL-MS data also implies conformational flexibility within ZUP1.

### The ZUP1 α-2/3 region promotes the formation of the ZUP1-RPA complex

We next sought to identify the RPA binding region(s) in ZUP1 by generating a range of GST-tagged ZUP1 fragments, whereby single or multiple domains were removed (Figure 4A). To ensure that the catalytic domain was functional in these assays, and therefore likely folded, truncated and full-length ZUP1 proteins were reacted with Ub-PA, which produced slower migrating species when analysed by SDS-PAGE (Extended Data Figure 4A). Using these ZUP1 truncations, we found that the N-terminal region (aa 1-269) of ZUP1, containing the three ZnFs, UBZ, MIU and ZHA, was insufficient to pulldown RPA (Figure 4B, lanes 10-12). In contrast, all ZUP1 fragments containing the *α*-2/3 and C78 peptidase domain were competent for RPA pulldown (Figure 4B, lanes 5-9). Removal of the *α*-2/3 region immediately N-terminal to the C78 peptidase domain abrogated the binding to RPA, suggesting that the ZUP1 *α*-2/3 region is primarily responsible for RPA binding *in vitro* (Figure 4B, compare lanes 3 and 4). This was the case for both truncated proteins lacking the *α*-2/3 region and in ZUP1 proteins with deletion of this domain alone, either with or without a short linker (Figure 4C and Extended Data Figure 4B). Importantly, we found that, whilst not completely abrogated, loss of the *α*-2/3 region, either in a truncated ZUP1 or by deletion of the domain alone, severely compromised the ability of FLAG-ZUP1 to co-immunopurify RPA from cells (Figure 4D). Given that the RPA70 N-terminal or RPA32 C-terminal interacts with numerous client proteins, we analysed the *α*-2/3 region for similarity to known RPA-interacting sequences. However, there was no obvious alignment of the *α*-2/3 region to known motifs of RPA70 or RPA32-interacting proteins, suggesting that the interaction interface remains elusive (Extended Data Figure 4C,D). Regardless, together, our biochemical and cellular data identifies the ZUP1 *α*-2/3 region as the primary domain responsible for promoting formation of the ZUP1-RPA complex (Figure 4E).

**Figure 4.**
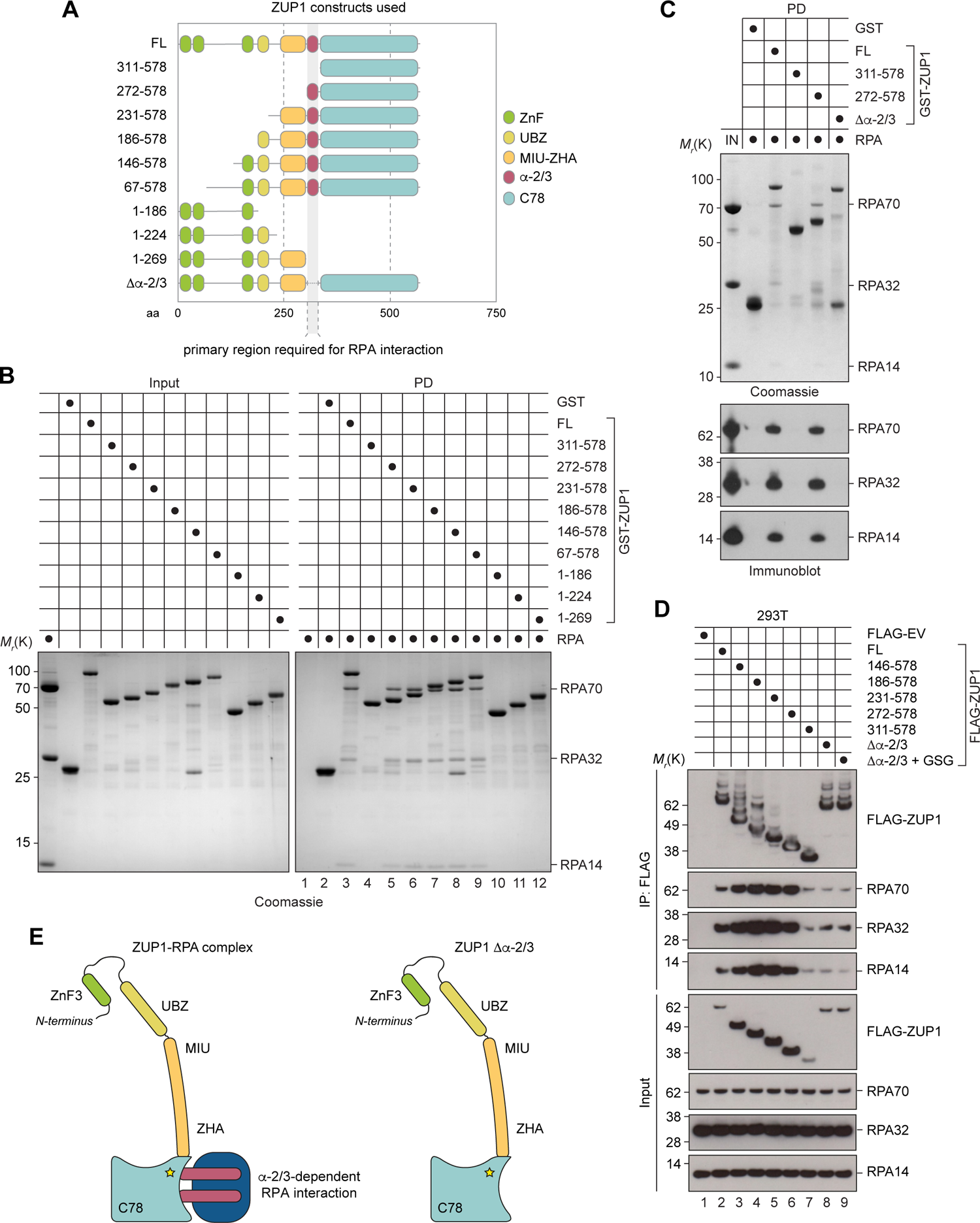
The ZUP1 *α*-2/3 region promotes the formation of the ZUP1-RPA complex. **A.** Domain diagram of full-length ZUP1 and the various mutants used in this paper. **B.** GST pulldowns using GST, full-length (FL) GST-ZUP1, or GST-ZUP1 truncated proteins, together with RPA. 20% of the input (IN) and 50% of the eluted pulldown (PD) was analysed by Coomassie staining. **C.** As in (B), but with additional immunoblotting. 10% of the input (IN) and 50% of the eluted pulldown (PD) was used for both the Coomassie-stained gel and for immunoblotting. **D.** 293T cells were transfected with FLAG-EV, FLAG-ZUP1 FL or the indicated truncation or deletion mutants, followed by FLAG immunoprecipitation (IP), and then analysed by immunoblotting with the indicated antibodies. Data is representative from at least *n* = 3 biological repeats. **E.** Model summarising the data in this figure. *Left*, the ZUP1 *α*-2/3 region is primarily responsible for the RPA interaction. *Right*, loss of the *α*-2/3 abrogates ZUP1-RPA complex formation. See also Extended Data Figure 4.

### ZUP1, RPA, and polyUb form a complex

Given that ZUP1 has previously been shown to bind to polyUb chains and we have shown here that ZUP1 can form a stable complex with RPA in the presence or absence of ssDNA, it raised the possibility that ZUP1 might bind to RPA and polyUb chains simultaneously. To address this, we performed gel filtration using a Superdex 200 column to analyse complex formation between a catalytically inactive mutant of full length ZUP1 (C360A), RPA, and K63-linked tetra-Ub (Ub4). Ubiquitin has poor absorbance at 280 nm, however, absorbance measurements at 214 nm allowed detection of complexes containing K63-linked Ub4 chains, with peak shifts detected for the ZUP1-RPA complex and a shift with a corresponding peak height increase for the ZUP1-RPA-Ub4 complex (Figure 5A). This SEC analysis therefore suggested that the ZUP1-RPA complex can also simultaneously bind Ub4. To extend this further, we used the XL-MS with SCX chromatography workflow described above to analyse the ZUP1-RPA-Ub4 complex. Crosslinks between ZUP1 and RPA were similar with or without K63-linked Ub4 chains (compare Figure 3B and Figure 5B,C), whilst the majority of crosslinks from Ub formed with residues in ZUP1 (Figure 5C,D). Crosslinks between lysine residues on Ub to K356 on ZUP1 place part of the Ub chain within the active site, while other regions within ZUP1 indicate Ub binding regions or regions of ZUP1 in close proximity to the polyUb chain bound to ZUP1, such as the N-terminal ZnFs, ZnF3 close to the UBZ, and the MIU (Figure 5D). We also detected crosslinks between RPA32 and RPA70 with ZUP1 K356 only in the presence of polyUb (Figure 5C). Given the repetitive nature of the K63-linked Ub4 chains, it is difficult at present to definitively locate the polyUb chain within the complex. However, despite this, collectively our XL-MS data reinforces our SEC analysis and indicates that ZUP1, RPA, and ubiquitin can form a complex *in vitro*.

**Figure 5.**
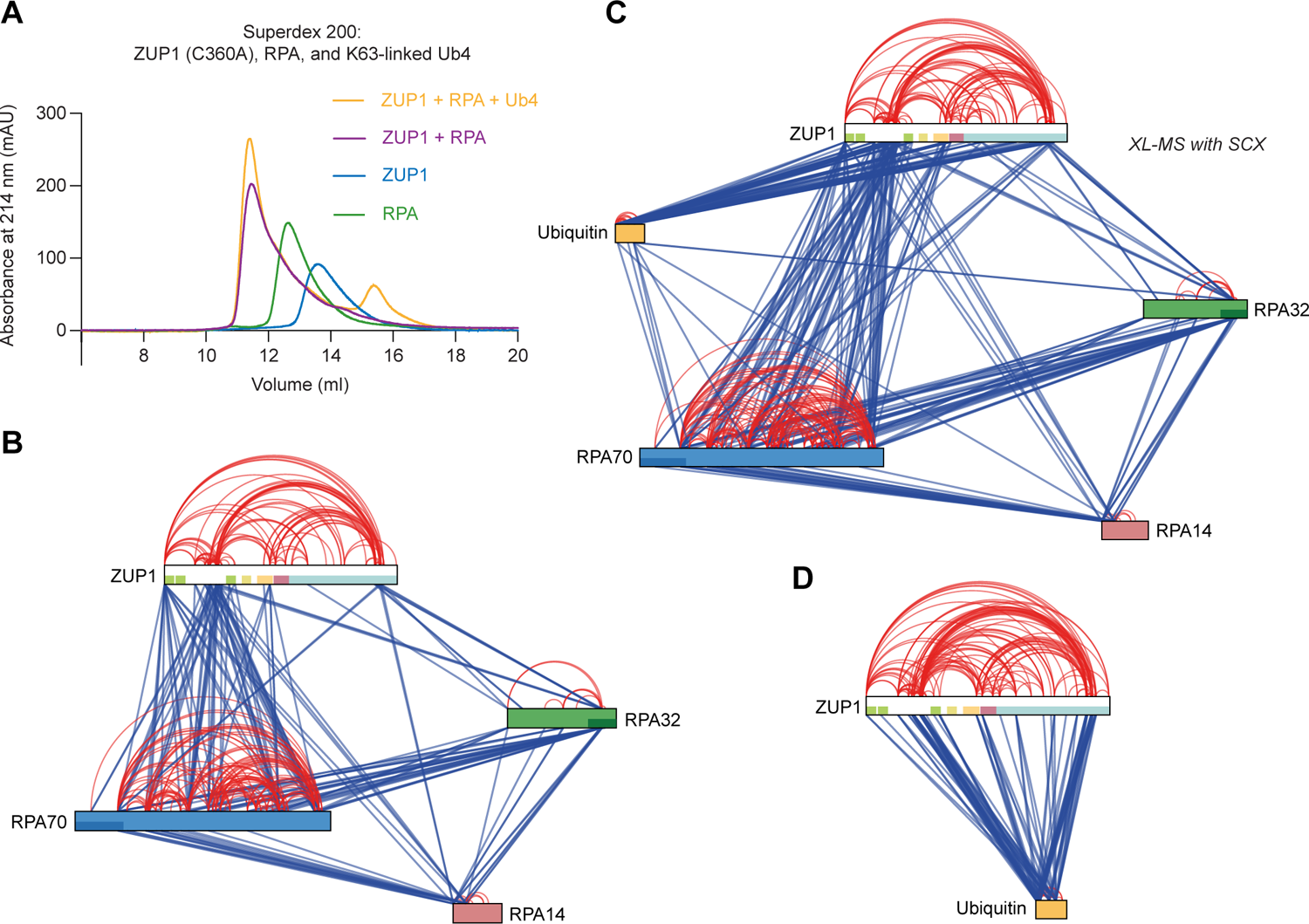
ZUP1, RPA, and ubiquitin form a complex. **A.** Chromatogram from analytical SEC showing the absorbance at 214 nm of the indicated proteins and protein complexes run over a Superdex 200 10/300 gel filtration column. Note that 214 nm was used to enable detection of Ub chains, which absorb poorly at 280 nm. Peak shifts can be detected when ZUP1 C360A (catalytically inactive point mutant) is pre-incubated with RPA and when ZUP1 C360A is pre-incubated with both RPA and K63-linked tetra-Ub (Ub4) chains. **B.** Same data as Figure 3B, for ease of comparison. **C.** As in (B), except with ZUP1 C360A and K63-linked Ub4 chains. **D.** As in (B), except with and ZUP1, RPA and K63-linked Ub4 chains. Ub is shown as a single species despite existing as a Ub4 molecule in these assays.

### RPA stimulates ZUP1 K63-linked deubiquitination activity

Our finding that the *α*-2/3 region promotes formation of the ZUP1-RPA complex was of particular interest as the *α*-2/3 has previously been shown to also be required for ZUP1 deubiquitination activity against K63-linked polyUb, and presumed to be the location for the proximal S1’ Ub ^20, 22, 24^. Moreover, given our findings here that ZUP1 can form a complex with both RPA and K63-linked polyUb, it raised the possibility that RPA-binding to ZUP1 might modulate its ability to hydrolyse K63-linked polyUb chains. We had initially hypothesised that RPA might inhibit ZUP1 activity by blocking polyUb chain access to the *α*-2/3 region. However, surprisingly, addition of stoichiometric amounts of RPA to *in vitro* DUB assays revealed that RPA could greatly stimulate the deubiquitinating activity of ZUP1, enhancing its ability to hydrolyse K63-linked chains into shorter fragments (Figure 6A). Importantly, by examining the ZUP1 C360A mutant, we ensured there was no contaminating DUB in the RPA preparations (Figure 6A). The RPA-dependent stimulation of ZUP1 DUB activity did not alter ZUP1’s specificity for K63 linkages within polyUb (Extended Data Figure 5A), whilst analysis of K63-linked Ub chain length showed that the degree of RPA-dependent stimulation increased with the lengthening of the Ub chain (Figure 6B and Extended Data Figure 5B-E). Previous data showed that ZUP1 can bind K6, K48, and K63-linked polyUb chains ^20^. We therefore generated heterotypic branched K48-K63 polyUb chains of differing lengths and found that the RPA-dependent stimulation of ZUP1 DUB activity remained specific for the K63 linkages within these heterotypic polyUb chains (Extended Data Figure 5F). Inclusion of ssDNA or phospho-RPA32 mutants had negligible impact on the RPA-dependent stimulation of ZUP1 DUB activity (Extended Data Figure 5G,H). Taken together, this data shows that RPA promotes ZUP1 DUB activity against K63-linked polyUb chains.

**Figure 6.**
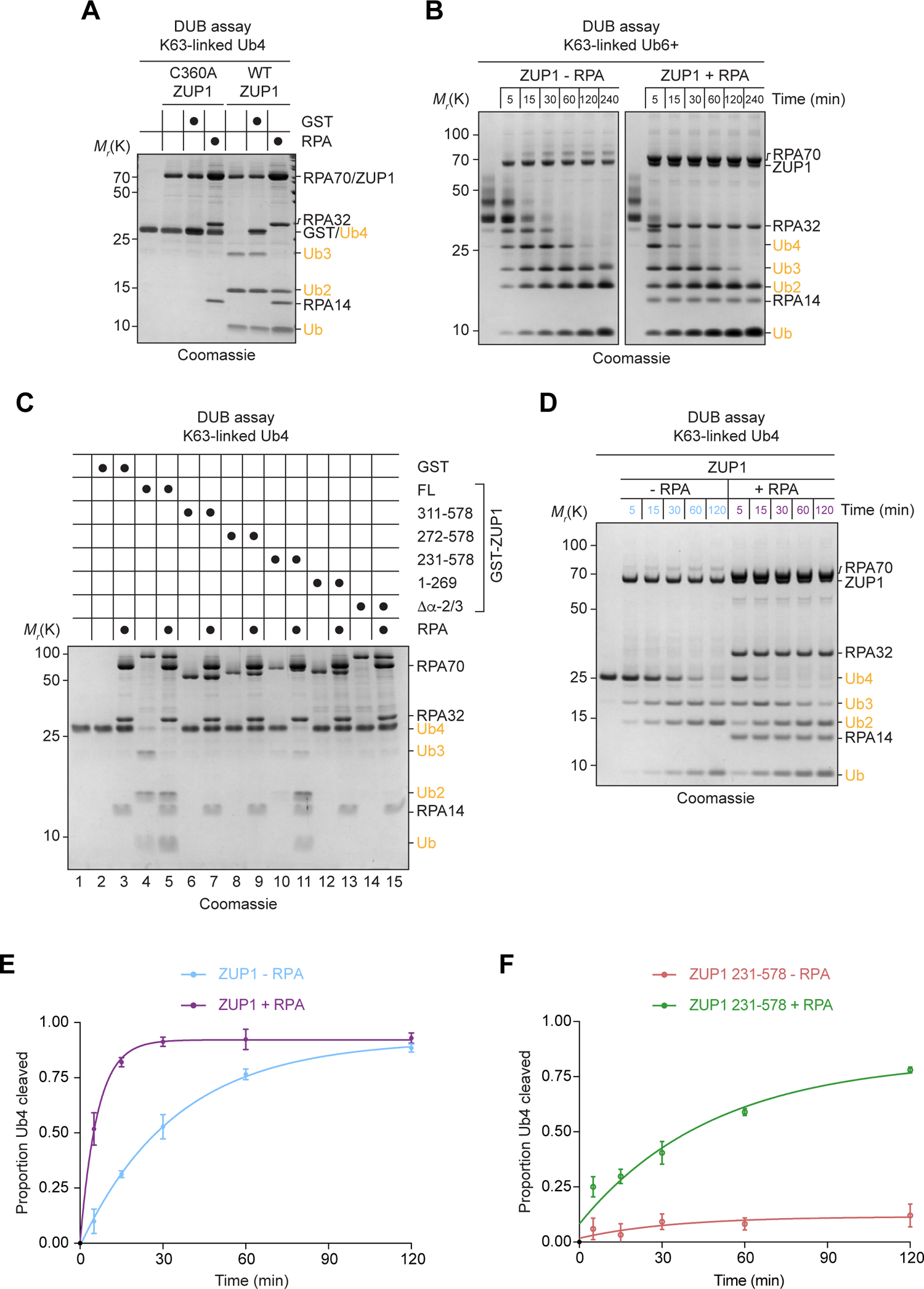
RPA stimulates ZUP1 K63-linked deubiquitination activity. **A.** *In vitro* DUB assay with the indicated following components: GST (1.25 µM), wildtype (WT) or C360A catalytically inactive ZUP1 (1 µM), K63-linked Ub4 chains (2 µM), and RPA (1.25 µM). The reaction was incubated for 2 h at 25°C and quenched by the addition of SDS loading buffer without boiling. The reaction products were separated SDS-PAGE and stained with Coomassie. **B.** As in (A), except K63-linked Ub6+ chains were used, with samples taken at the indicated time points. **C.** As in (A), except various ZUP1 truncation or deletion mutants were used. **D.** As in (A), except samples were taken at the indicated time points. **E.** Quantification of the Ub4 band from (D). Data is from *n* = 3 independent experiments and represents the mean and standard deviation. **F.** As in (E), except using ZUP1 231-578 protein. See also Extended Data Figure 5.

If the ZUP1 *α*-2/3 is indeed the S1’ site for ZUP1, then our data suggests a functional bifurcation of the *α*-2/3 region, whereby it is important for both RPA binding and Ub chain hydrolysis. We next sought to characterise this phenomenon in more detail. As expected, loss of the *α*-2/3 abrogated ZUP1 DUB activity, in the presence or absence of RPA (Figure 6C – compare lanes 4 and 5 with 6, 7, 14, and 15). Interestingly, though the ZUP1 *α*-2/3-C78 (aa 272-578) fragment interacted with RPA (Figure 4), the DUB activity of this fragment could not be stimulated by addition of RPA (Figure 6C – compare lanes 4 and 5, with 6-9). However, addition of the MIU-ZHA domain to this fragment (generating aa 231-578) was sufficient to reinstate RPA-dependent stimulation of ZUP1 (Figure 6C – compare lanes 4 and 5 with 10 and 11). Thus, whilst the UBZ domain is required for the maximum basal (i.e., in the absence of RPA) DUB activity against K63-linked polyUb chains, it is not strictly required for the RPA-mediated stimulation of ZUP1 DUB activity (Figure 6B). This data suggests that RPA stimulates ZUP1 DUB activity at the very least through the MIU-ZHA domain, a region which includes the S1 site and probable S2 site.

The surprising finding that the UBZ domain is not strictly required for the RPA-dependent stimulation of ZUP1 DUB activity prompted us to assess this further using DUB assays with greater temporal granularity. Using this approach, the impact of RPA on the rate of disappearance of Ub4 and appearance of Ub*≤*3 was strikingly evident (Figure 6D,E). Repeating this approach for ZUP1 231-578 showed that in the absence of the UBZ, ZUP1 has a severely compromised basal activity against K63-linked polyUb, as previously reported, yet the addition of RPA is sufficient to drive the reaction to near completion, albeit at a slower rate (Figure 6F). Furthermore, we could recapitulate these findings using a UBZ mutant (*UBZ) within the context of full-length ZUP1, rather than a truncation (Extended Data Figure 5I,J). Together, this data suggests that the *α*-2/3-dependent interaction with RPA promotes enzyme-substrate complex formation via the MIU-ZHA.

Based on the data above and given that the ZHA and *α*-2/3 are contiguous within the ZUP1 polypeptide chain, we postulated that RPA binding to ZUP1 might stimulate the ZHA S1 site in ZUP1. To address this, we performed DUB assays with Ub-AMC, a minimal DUB substrate in which the AMC fluorophore is conjugated to the C-terminus of Ub, with cleavage resulting in AMC fluorescence (Extended Data Figure 5K). Importantly, as the Ub moiety within the Ub-AMC binds to the distal S1 Ub binding site in DUBs, it gives a direct indication of the binding of the ZUP1 ZHA. Similar to the K63-linked polyUb chain DUB assays, addition of RPA stimulated ZUP1 DUB activity, with a 10-fold increase in the efficiency of Ub-AMC cleavage (k_cat_/K_M_) with RPA compared to a GST control (Extended Data Figure 5L-Q). The RPA-dependent stimulation of ZUP1 DUB activity was almost identical for both full-length ZUP1 and 231-578 fragment, indicating that the mechanism of stimulation against Ub-AMC is independent of the UBZ domain, as expected, and that RPA binding stimulates the ZHA S1 site (Extended Data Figure 5M,N,P,Q).

Collectively, our data suggests that RPA stimulates ZUP1 DUB activity exclusively against K63-linkages within homotypic and heterotypic polyUb chains. Mechanistically, we show that this RPA-dependent stimulation is mediated through at least the ZHA S1 site, suggesting that RPA binding exerts its stimulatory impact by promoting enzyme-substrate complex formation. Furthermore, whilst the UBZ is not strictly required for RPA-mediated stimulation of ZUP1 DUB activity against polyUb chains or a minimal substrate, it is still necessary for maximal ZUP1 activity against longer K63-linked polyUb chains together with RPA. Thus, both the RPA-dependent S1 site stimulation and the UBZ domain independently contribute to maximal ZUP1 DUB activity, though the former has the greater impact on ZUP1 activity.

### RPA binding is the main mechanism to stimulate ZUP1 deubiquitinase activity

The RPA-dependent stimulation of ZUP1 DUB activity raised the question of how specific this is to both ZUP1 and RPA. RPA showed no stimulation of USP2 or USP11 activity when assessed by *in vitro* DUB assays, indicating that RPA does not stimulate other DUBs (data not shown). In addition, the DNA replication and repair factor PCNA, also an abundant protein at DNA replication forks and target of ubiquitination like RPA, was unable to stimulate ZUP1 DUB activity, even when titrated far in excess of ZUP1, indicating that it is unable to fulfil the role of ZUP1 activator (Figure 7A and Extended Data Figure 6A). Whilst this is far from an exhaustive list of DUBs and DNA replication and repair factors, thus far we have observed that the stimulation of ZUP1 DUB activity is specific to RPA.

**Figure 7.**
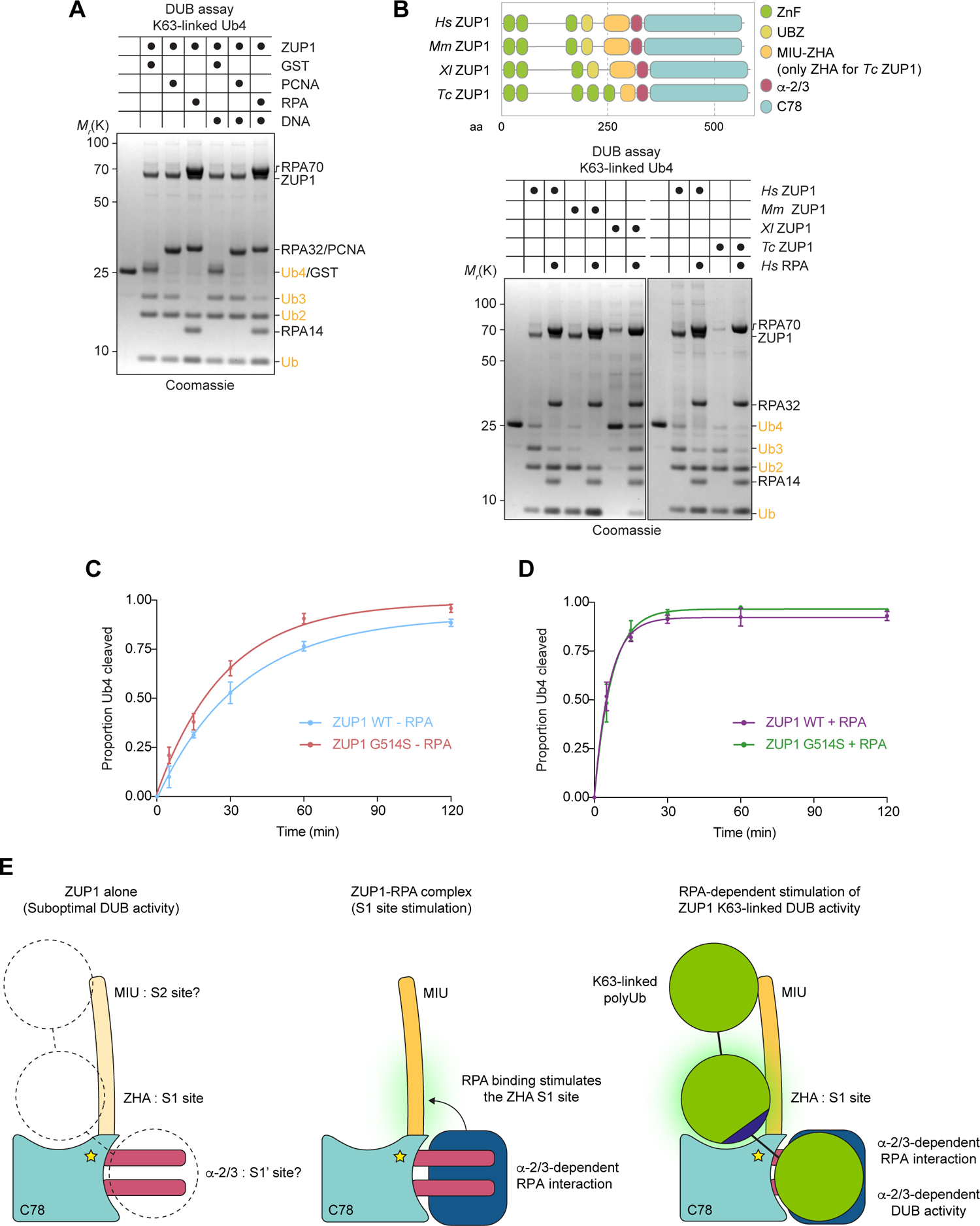
RPA is the main mechanism to stimulate ZUP1 deubiquitinase activity and is evolutionarily conserved. **A.** *In vitro* DUB assay with the indicated following components: GST (1.25 µM), ZUP1 (1 µM), K63-linked Ub4 chains (2 µM), RPA (1.25 µM), PCNA (1.25 µM), and ssDNA. The reaction was incubated for 2 h at 25°C and quenched by the addition of SDS loading buffer without boiling. The reaction products were separated SDS-PAGE and stained with Coomassie. **B.** As in (A), except with three additional ZUP1 homologs, mouse (*Mus musculus*, *Mm*ZUP1; 1.25 µM), frog (*Xenopus laevis*, *Xl*ZUP1; 1 µM), and insect (*Tribolium castaneum*, *Tc*ZUP1; 0.2 µM). **C.** Quantification of the Ub4 band from *in vitro* DUB assays containing ZUP1 WT or G514S, together with K63-linked Ub4 chains. **D.** As in (D), except RPA was added to the reaction. **E.** Model for RPA-dependent stimulation of ZUP1 DUB activity. *Left*, in the absence of RPA, ZUP1 is reliant on the UBZ (not shown) for basal activity, yet even when the UBZ is present ZUP1 functions at suboptimal rates on K63-linked polyUb substrates. Putative S1’ and S2 sites are shown. Star indicates the ZUP1 active site in the C78 domain. *Middle*, the ZUP1 *α*-2/3 promotes ZUP1-RPA complex formation, which in turn stimulates the ZUP1 ZHA S1 site. *Right*, binding of RPA stimulates enzyme-substrate formation. The *α*-2/3 is also required for ZUP1 DUB activity in the absence of RPA, suggesting it is the likely S1’ site. Blue indicates R-x-R motif in Ub. See also Extended Data Figure 6.

Given that ZUP1 is generally well conserved in eukaryotes, we expressed and purified three other homologs and analysed whether human RPA was sufficient to stimulate their DUB activity ^20–22, 24^. We chose mouse (*Mus musculus*, *Mm*ZUP1) and frog (*Xenopus laevis*, *Xl*ZUP1) ZUP1 homologs as they share the exact same domain architecture with human (*Hs*ZUP1), with AlphaFold predictions also suggesting that these are all structurally similar proteins (Extended Data Figure 6B). We also chose insect ZUP1 (*Tribolium castaneum*, *Tc*ZUP1) as a third homolog as it contains an additional ZnF in place of the MIU domain equivalent in *Hs*ZUP1 (Figure 7B and Extended Data Figure 6B). Notably, all three homologs contain the ZHA and *α*-2/3. Whilst the basal DUB activities of the homologs varied somewhat, we observed clear RPA-dependent stimulation of DUB activity across all three homologs, suggesting that this mechanism is likely conserved in species with at least the ZHA, *α*-2/3 and C78 domains (Figure 7B).

As we have shown above, when ZUP1 is present within the ZUP1-RPA complex, the role of the UBZ becomes less important for ZUP1 DUB activity than when ZUP1 is alone. From this we concluded that the RPA-dependent stimulation is the major mechanism of ZUP1 activation. Recently, whilst trying to understand how the oxyanion hole contributes to ZUP1 activity in different homologs, the Hofmann lab identified *Hs*ZUP1 Gly514 as a residue whose mutation to serine, its equivalent in *Tc*ZUP1, could modestly increase the activity of *Hs*ZUP1 against K63-linked polyUb (6+) chains ^24^. We therefore made the same G514S mutation in *Hs*ZUP1 and found a similar modest increase in activity against K63-linked Ub4 chains (Figure 7C,D and Extended Data Figure 6C,D). However, when RPA was included in the assay, the *Hs*ZUP1 G514S mutant made little difference to the DUB activity versus *Hs*ZUP1 wild type (Figure 7C,D and Extended Data Figure 6C,D). Taken together with the data above, this suggests that the RPA-dependent stimulation is the main activating mechanism for ZUP1 DUB activity in species whose ZUP1 gene product contains the ZHA, *α*-2/3 and C78 domains.

## Discussion

While ZUP1 was previously shown to be a K63-sepcific DUB, its activity is very poor, raising the question of how it exerts its biological function. In this study, we provide novel insights into the biochemical basis of ZUP1 DUB activity. We show that ZUP1 directly interacts with the major ssDNA binding protein in eukaryotes, the RPA complex, both in cells and *in vitro*. Previous models of ZUP1’s relationship to RPA posited that it may be indirect, and that some unknown factor(s) might mediate their interaction. However, our data suggests a simpler mechanism that couples ssDNA detection to K63-linkage specific deubiquitination via the formation of the ZUP1-RPA complex. We find that the ZUP1 *α*-2/3 region, the potential S1’ site in ZUP1, is important for ZUP1-RPA complex formation, though other interaction sites are likely based on our XL-MS data, suggesting a more complex mode of binding compared to other RPA-interacting partners such as SMARCAL1 and p53 (Extended Data Figure 4D). This is reminiscent of the E3 ligase RAD18, which has been shown to bind to multiple subunits within RPA and its activity is regulated by the interaction with RPA ^31, 45, 46^. Importantly, we show that RPA functions to stimulate the DUB activity of ZUP1 against K63-linked polyUb chains. Mechanistically, we show that RPA stimulates ZUP1 DUB activity through interdomain communication between the S1’ and S1 sites within ZUP1, which functions in parallel to the UBZ domain to optimise overall ZUP1 DUB activity (Figure 7E).

Taken together, our results imply that the ZUP1-RPA complex deubiquitinates substrates modified with K63-linkages in long polyUb chains at ssDNA regions (Extended Data Figure 7). There is growing evidence that RPA coordinates Ub signaling at ssDNA via interaction with a number of Ub E3 ligases, such as RAD18 ^31^, RFWD3 ^32–35^, PRP19 ^36^, and HERC2 ^37^. Some of these E3 ligases have been shown to ubiquitinate RPA, with various functions suggested. For PRP19, it was proposed to promote the activation the ATR kinase ^36^, whilst for RFWD3 it has been suggested to promote homologous recombination (HR) at stalled forks, by preventing inappropriate retention of RPA on ssDNA and allowing downstream RAD51 loading ^33–35^. Alternatively, it was recently shown that RFWD3 ubiquitinates proteins on ssDNA, thereby recruiting numerous factors in a Ub-dependent manner, including proteins involved in promoting PCNA ubiquitination ^32^. Consequently, the absence of RFWD3 leads to severe defects in TLS at ssDNA gaps. Whether ZUP1 counteracts any of the processes mediated by these RPA-E3s will need to be examined, as well as whether RPA orchestrates deubiquitination of other Ub linkages via different DUBs. Lastly, as ssDNA gaps have recently been proposed to underlie the PARPi sensitivity of *BRCA1/BRCA2*-mutated cells, it will be also important to determine if ZUP1 has a function in these pathways ^47, 48^.

Here we identify RPA as a cofactor that activates ZUP1, which adds it to a growing list of DUBs which require cofactors for activation. For example, Ubp8 (USP22 in humans) in the SAGA complex, requires four additional proteins, Sgf11, Sus1, and Sgf73, that form into a subcomplex for its ubiquitin binding and activity ^49^. A second prominent example is USP1, a DUB involved in removing the monoUb from PCNA and the FANCD2-FANCI complex and which requires its cofactor UAF1 to enhance its DUB activity and substrate specificity ^50, 51^. We show here that the ZUP1 *α*-2/3, the likely S1’ site, promotes interaction with RPA, both *in vitro* and in cells, and in doing so stimulates enzyme-substrate formation at the ZHA S1 site. This was both surprising and of particular interest as the *α*-2/3 has been previously suggested to be the S1’ site in ZUP1 ^20, 22, 24^. What is the evidence that the *α*-2/3 might be S1’ site? Firstly, from the two structures of human ZUP1 bound to Ub-PA, the ZHA S1 site positions the C-terminal of the distal Ub towards the *α*-2/3, where the proximal Ub and K63-linkage would be found ^20, 22^. Secondly, as shown here and by the Hofmann group ^22, 24^, deletion of the *α*-2/3 almost completely abolishes ZUP1 activity, underlining its critical role in catalysis. At present, exactly how the *α*-2/3 contributes to ZUP1 K63-linkage specificity remains elusive. However, as we have shown here, the K63-linkage specificity of ZUP1 is independent of the ZUP1-RPA interaction, thus the K63-linkage specificity must reside within the ZUP1 polypeptide. Previous attempts to disrupt the *α*-2/3 through multisite mutation didn’t alter the properties of ZUP1, so it was suggested that its ability to bind Ub as the S1’ site might be mediated via main chain interactions ^24^. Another possibility is that the ZUP1 C78 peptidase domain contributes some interacting residues to the S1’ site that, by themselves, are not sufficient for robust interaction between the C78 domain and ubiquitin, which previous studies couldn’t detect ^20–23^. In contrast to the suggestion that the *α*-2/3 acts solely as the S1’ site, it is also possible that the ZUP1 UBZ acts, or at least contributes to, the S1’ site, perhaps together with the *α*-2/3 ^20^. The use of UBDs in the S1’ site, which can occur in *cis* or in *trans*, is a known mechanism for promoting DUB activity ^15^. Examples include the UBP-ZnF domain in USP5 (IsoT), the AnkUBD in TRABID, and the UIM domain in OTUD1 ^52–54^. Introduction of inactivating point mutants in the UBZ domain, either in truncated or full-length ZUP1, compromise ZUP1 activity against polyUb chains, as we have shown here and previously ^20^. For this to be possible, there would likely need to be some local unfolding of the MIU to allow this orientation (Extended Data Figure 1E). Our XL-MS supports this idea of conformational plasticity in ZUP1, as we observe both long-range crosslinks between the N- and C-terminus and local crosslinks greater than the distance expected of the BS3 crosslinker. It remains to be determined whether the MIU-ZHA is structured in solution in the absence of polyUb binding. Furthermore, it is noteworthy to point to the predicted structure of *Tc*ZUP1, with its ZnF domains connected by flexible linkers, which might easily contribute to the S1’ site (Extended Data Figure 6B). In addition, in plants, the *α*-2/3 is replaced by a UBZ, which takes on the role of the S1’ site ^24^. Collectively, whilst the mechanisms of K63-linkage specificity and polyUb chain recognition remain to be determined for ZUP1, we now have the basis to uncover these using ensemble approaches in light of its newly identified activating partner, RPA.

As noted above, we have yet to determine the structural basis of the ZUP-RPA-Ub4 complex, which therefore remains an important future challenge. Thus, we can only speculate as to the structural basis for the RPA-dependent stimulation of ZUP1. However, there are some important points from our data that may point towards how RPA might achieve this. Firstly, that RPA activates, rather than inhibits, ZUP1 activity suggests that RPA doesn’t occupy the S1’ site itself, as this site will present the Ub K63-linkage to the catalytic site. Secondly, the SEC data suggests that ZUP1 C360A-RPA-Ub4 can form a complex. Though the SEC data does not formally rule out that RPA occupies the S1’ site, it seems incongruous with the stimulation of ZUP1 DUB activity if it did. Thus, we have concluded that there is likely a functional bifurcation of the *α*-2/3 region. That is, it functions to link ZUP1 to RPA, whilst at the same time acts as part of the S1’ site. A reasonable suggestion for how it achieves this would be a spatial separation of the two functions, with the proximal Ub binding on the ‘top’ and RPA binding on the ‘side’ or ‘bottom’ of the *α*-2/3 region. Whilst this may explain the two functions of the *α*-2/3 region, it does not explain how RPA stimulates ZUP1 activity and connects binding at the S1’ site to S1 site activation. Perhaps the simplest explanation is that the RPA complex allows better presentation of the K63-linkage to the active site via improved orientation of the ZHA-*α*-2/3 relative to the C78 domain. This might occur if RPA physically connects the S1 and S1’ sites. Whilst we have shown that the *α*-2/3 is necessary for the RPA interaction, we have been unable to show whether it is sufficient, despite attempts to produce the isolated *α*-2/3. Alternatively, if RPA does not physically connect the S1 and S1’ sites, then given that the *α*-2/3 region is contiguous with the ZHA, RPA binding may help induce secondary structure upstream of the *α*-2/3, potentially from the end of the ZHA to the start of the MIU, as this is one long *α−*helix from published structures and AlphaFold predictions. In this scenario, RPA might help position the *α*-2/3 into a conformation that better presents the K63-linkage to the active site, whilst also perhaps limiting the ability of the ZHA and MIU to become unstructured. Further approaches to understand the dynamics of the MIU-ZHA with and without Ub binding should help clarify these, in addition to determining the structural basis of the ZUP1-RPA-Ub complex.

We show here that the RPA-dependent stimulation of ZUP1 DUB activity functions in parallel to the UBZ for optimal ZUP1 polyUb chain hydrolysis. The UBZ domain, as the principle Ub-binding domain in ZUP1 ^20, 22^, might assume at least two locations depending on the presence or absence of the RPA. Firstly, it may contribute to the S1’ site in conjunction with the *α*-2/3 as mentioned above, or it might interact with distal units in the polyUb chain in the S3 site (Extended Data Figure 1D,E). It will be important to determine which of these orientations is possible for ZUP1 alone and within the ZUP1-RPA complex. It is possible that the UBZ might be key for ZUP1 to ‘read’ the length of the Ub chain, and as it does so co-operate with RPA to help form the long MIU-ZHA alpha helix. Understanding the affinities of individual UBDs in ZUP1 together with single-molecule approaches will help determine if such a mechanism exists.

The ZUP1-RPA complex seems to be constitutive in cells under all the conditions we have analysed. Therefore, it seems likely that the RPA interaction might also contribute to its recruitment to ssDNA. Indeed, the ZUP1-RPA complex forms on all ssDNA-containing substrates we have examined *in vitro*. As such, the ZUP1-RPA complex may be primed to hydrolyse long K63-linked polyUb chains from substrates on or near ssDNA. As we have shown here, the K63-linkage could be within homotypic or heterotypic polyUb chains. There are several implications of this and questions to address. Firstly, are there are other PTMs that further regulate the ZUP1-RPA complex in response to DNA damage, particularly ZUP1 itself, having shown here that RPA phosphorylation doesn’t impact ZUP1 activation? Secondly, what are the substrates upon which ZUP1 acts? At present, there is only one suggested substrate of ZUP1, the RPA complex itself ^23^, however the functional consequences of cellular hyper RPA polyUb in the absence of ZUP1 remains to be determined. It seems unlikely to us that RPA is the sole substrate of ZUP1, given the range of E3 ligases at ssDNA and the subsequent requirement for a mechanism to strictly limit ZUP1 activity to target RPA polyUb. Thus, it will be imperative to map the substrates of ZUP1 using Ub proteomic tools, as well as the polyUb chain architectures on these substrates ^55^. Thirdly, it is likely that there are more components to the ZUP1-RPA complex in a cellular context. The identity of these components and how they contribute to the recruitment, activity, and regulation of the ZUP1-RPA complex at DNA lesions will also need to be investigated. Fourthly, if the ZUP1-RPA complex is an active DUB against K63-linked polyUb chains at ssDNA, then how will ZUP1 activity be negatively regulated to ensure sufficient signaling capacity of the K63-linked polyUb chains? Lastly, the answers to these questions will need to be framed within the context of which DNA repair pathway(s) ZUP1 functions within, which remains to be fully defined.

The RPA-dependent stimulation of ZUP1 activity was evident in three additional ZUP1 homologs, suggesting that the mechanism is evolutionarily conserved. From these observations, and considering the collective findings presented here, we infer that the specific arrangement of domains involved in this mechanism have evolved to promote RPA-dependent stimulation of ZUP1 DUB activity. The three minimal domains required are the ZHA (S1 site), *α*-2/3 (RPA-binding/S1’ site), and C78 domains (catalytic activity). Given that these same three domains are also required for the K63-linkage specificity and minimal activity of ZUP1 (not the full activity – this requires the additional UBDs), it will be interesting to determine whether additional species which contain this domain arrangement have maintained the RPA-dependent stimulation. ZUP1 homologs are present in many eukaryotic supergroups and whilst it is notable that the domains upstream of the C78 domain vary considerably depending on the supergroup (Extended Data Figure 8A), the ZHA-*α*-2/3 domain arrangement is absent in Holomycota and only present in metazoa (Extended Data Figure 8B,C). The emergence of the ZHA-*α*-2/3 domain arrangement in metazoa suggests that it coincided with a functional requirement for RPA-mediated K63-linked deubiquitination in DNA repair pathways.

Overall, our findings identify a mechanism by which the recently discovered ZUP1 DUB is activated in a context-dependent manner by the RPA complex. Our discoveries therefore suggest a mechanism that couples the detection of ssDNA to the activation of K63-linked deubiquitination via the ZUP1-RPA complex.

## Acknowledgements

I.G.-S. gratefully acknowledges support from a Cancer Research UK Career Development Fellowship (C62538/A24670), the Oxford University Press John Fell Research Fund (0006091), the University of Oxford Medical Sciences Division Pump Priming Award (006353), and the Brasenose College Research Fund. K.C. is supported by an Elizabeth Murphy Scholarship and M.D. is supported by a BBSRC Doctoral Training Programme studentship, both from the University of Oxford. We thank Francis Barr (Department of Biochemistry, University of Oxford, UK), Paul Elliott (Department of Biochemistry, University of Oxford, UK), Ivan Ahel (Sir William Dunn School of Pathology, University of Oxford, UK), John Rouse (MRC PPU, University of Dundee, UK), and Marc Wold (Department of Biochemistry and Molecular Biology, University of Iowa, USA) for providing reagents. We sincerely thank Yogesh Kulathu (MRC PPU, University of Dundee, UK) for critical reading of the manuscript.

## Author Contributions

Conceptualization, B.F. and I.G.-S.; Methodology, B.F., K.C., and M.F.; Formal analysis, B.F. and I.G.-S.; Investigation, B.F., M.A., K.C., M.D., R.G. I.G.-S.; Resources, M.F.; Data curation, M.F.; Writing – Original Draft, B.F. and I.G.-S.; Writing – Review and Editing, B.F., M.A., I.G.-S.; Visualization, B.F. and I.G.-S.; Supervision, B.F. and I.G.-S.; Project administration, I.G.-S.; Funding acquisition, I.G.-S.

## Materials and Methods

### Cell culture

Human HEK293T (referred to as 293T) were purchased from ATCC and cultured in DMEM (Thermo Fisher) supplemented with 10% fetal bovine serum (Thermo Fisher). Cells were grown in a 5% CO2 incubator at 37°C. Cells were routinely tested for mycoplasma contamination. For drug treatments, media was replaced with fresh DMEM containing either vehicle (DMSO) or the appropriate compound at the following concentrations: 5 µM ATRi (VE-822), 1 µM MMC, 1 µM Ub E1i (TAK-243), 1 µM SUMO E1i (ML-792). Plasmid transfections were performed with Opti-MEM (Thermo Fisher) mixed with sequence-verified plasmid DNA and PEI MAX transfection reagent (Polysciences) at a 1:3 mass ratio of DNA:PEI. Typically, 7.5 µg DNA was used per 10 cm dish and mixed with 1.5 ml Opti-MEM, incubated for 30 min, and added dropwise to cells. Cells were harvested 24-48 h post-transfection.

### Immunoprecipitation and immunoblotting

Cells were transfected with pcDNA5-FRT/TO-FLAG, pcDNA5-FRT/TO-FLAG-ZUP1, or pcDNA5-FRT/TO-RFWD3. After 48 h, cells were washed with ice-cold PBS and subsequently lysed with TX-100 buffer (50 mM Tris-HCl pH 8.0, 100 mM NaCl, 1% Triton X-100) complete with additives (>500 units/ml benzonase, 10 mM β-glycerophosphate, 1 mM DTT, 1 mM NaF, 5 mM NEM, 0.1 mM sodium orthovanadate, protease inhibitor cocktail (Sigma), 1 μM KU-55933 (ATMi), 1 μM NU7441 (DNA-PKi), 1 μM Olaparib (PARPi), 1 μM VE-822 (ATRi)).

Lysates were rotated for 20 minutes at 4°C and centrifuged at 13,000 g for 10 min at 4°C before discarding the pellet. From the clarified protein lysates, the protein concentrations of samples were measured against protease-free BSA standards using the Bio-Rad Quick Start Protein Assay protocol. Clarified lysates were normalised across samples and were added to 100 μl anti-FLAG M2 affinity gel (Merck) Micro Bio-Spin columns (Bio-Rad) pre-washed with 3x column volume of TX-100 buffer. Columns were rotated for 15 min at 4°C. Unbound lysate was drained by gravity flow and discarded. The FLAG co-immunoprecipitated cell lysates were washed five times with TX-100 buffer, before eluting with 1X LDS sample buffer containing 0.1 M DTT. Samples were heated, subjected to SDS-PAGE and analysed by immunoblotting. The following primary antibodies were used: mouse monoclonal ZUP1 (generated here), mouse monoclonal RPA32 (Abcam, ab2175), mouse monoclonal Ub (P4D1) (Thermo Fisher, 14-6078-82), mouse monoclonal SUMO antibody (8A2) (Abcam, ab81371), mouse monoclonal FLAG-HRP (Sigma-Aldrich (Merck), A8592), mouse monoclonal Vinculin-HRP (Santa Cruz Biotechnology, sc-73614-HRP), rabbit polyclonal RPA14 (Novus Biologicals, NBP1-87141), rabbit polyclonal RPA32 (Bethyl, A300-244A), rabbit polyclonal RPA70 (Cell Signaling Technologies, 2267S), rabbit monoclonal RPA70 (Abcam, ab79398), rabbit polyclonal Ub (Cell Signaling Technologies, 3933), rabbit monoclonal K63-specific Ub (Apu3) (Merck, 05-1308), and rabbit monoclonal K48-specific Ub (Apu2) (Merck, 05-1307). The following secondary antibodies were used: donkey anti-mouse IgG HRP-conjugated (Jackson ImmunoResearch, 715-035-150-JIR), donkey anti-rabbit IgG HRP-conjugated (Jackson ImmunoResearch, 711-035-152)

### Molecular biology

All plasmids were generated by using some combination of PCR amplification using Q5 DNA polymerase (NEB), HiFi Assembly (NEB, as per manufacturer’s instructions), and linearisation using HindIII and KpnI restriction enzyme digestion followed by annealing. Site-directed mutagenesis to introduce insertions, deletions, or amino acid substitutions was carried out using the Q5 DNA polymerase and KLD enzyme mix (NEB). Plasmid DNA was extracted and purified from 5 ml or 250 ml bacterial culture using Miniprep (Qiagen) or Maxiprep (Thermo-Scientific) kits, respectively, following the manufacturer’s instructions. The DNA concentration was measured using a Nanodrop spectrophotometer and verified by restriction digest and Sanger sequencing.

### Purification of recombinant proteins

For ZUP1-containing constructs, transformed *E. coli* Rosetta2 (DE3)/pLysS were grown to OD_600_ 0.6 in LB broth and then induced overnight at 18°C with 0.2 mM IPTG in media supplemented with 0.2 mM ZnCl_2_. Cells were harvested by centrifugation and re-suspended in lysis buffer (20 mM Tris pH 7.5, 500 mM NaCl, 20 mM imidazole, 0.1% Triton X-100, and 1x EDTA-free protease inhibitors (Roche)) and stored at −80°C until use. Cells were lysed by sonication on ice and clarified by centrifugation at 21,000 x *g* for 30 min at 4°C. Cell lysates were filtered using a 0.45 µM PES filter and incubated with Ni-NTA beads (Thermo Scientific), then equilibrated in lysis buffer without the protease inhibitors for 2 h at 4°C. Beads were sequentially washed with high salt (20 mM Tris pH 7.5, 500 mM NaCl, 20 mM imidazole) and low salt wash buffer (20 mM Tris pH 7.5, 100 mM NaCl, 20 mM imidazole) before re-suspension in low salt wash buffer. His-SENP1 (made in-house) was added to the re-suspended beads and incubated overnight at 4°C with rotation. The subsequent flow-through containing untagged ZUP1 was loaded onto a HiTrap Q HP 1 ml column (Cytiva) equilibrated in 20 mM Tris pH 7.5, 100 mM NaCl, 1 mM DTT and untagged ZUP1 was eluted using linear gradient from 100 mM to 1 M NaCl in 20 mM Tris pH 7.5, 1 mM DTT over 20 column volumes. Peak fractions were analysed by SDS-PAGE and Coomassie staining and fractions containing ZUP1 were pooled, concentrated using 10 kDa MWCO spin concentrators (Amicon) and loaded onto a Superdex 200 size-exclusion chromatography (SEC) column (Cytiva). ZUP1 was eluted by isocratic elution in 20 mM Tris pH 7.5, 100 mM NaCl, 1 mM DTT. Pure fractions (>90% by Coomassie Brilliant Blue-stained 10% SDS-PAGE) were pooled, concentrated and glycerol was added to 10% (v/v) before storage at −20°C (short-term) or −80°C (long-term). Full-length ZUP1 cDNA from mouse, frog, and insect homologs were ordered as synthetic genes from Thermo Fisher, then cloned, expressed, and purified in an equivalent manner. Other untagged ZUP1 mutants were generally purified in an equivalent manner except for 231-578 (MIU-C78), which used a HiTrap SP HP 1 ml column (Cytiva) due to an altered isoelectric point (pI). For GST-tagged ZUP1 proteins, expression was carried out in an equivalent manner as above. For purification, similar buffers were used except for the omission of imidazole in wash buffers and Glutathione Sepharose 4B beads (Cytiva) were used during the affinity purification step. The GST-tagged proteins were eluted using wash buffer supplemented with 20 mM reduced glutathione at pH 7.5. The eluate was concentrated and further purified by SEC as above. ZUP1 protein was purified from bacteria >4 times and all protein preparations produced equivalent yields and activity in experimental assays.

Human RPA complex was expressed from a pRSFDuet-1 vector with RPA70 inserted into MCS1 with an N-terminal His_6_-tag (with a short linker of SQDPNSSS) and RPA14 and RPA32 inserted in tandem into MCS2 without tags with separate ribosome binding sites (RBS) for each. Single colonies of transformed *E. coli* Rosetta2 (DE3)/pLysS were picked and directly added to 1 L of LB broth supplemented with kanamycin and chloramphenicol and incubated overnight at 25°C without shaking. The following day, the culture was grown at 37°C with shaking until reaching OD_600_ 0.6 and induced with 0.5 mM IPTG for 3 h at 37°C. Cells were harvested by centrifugation and re-suspended in J0 buffer (30 mM HEPES pH 7.5, 0.5% myo-inositol, 0.02% Tween-20) supplemented with 500 mM NaCl and 1x EDTA-free protease inhibitors (Roche) and stored at −20°C until use. Cells were lysed by sonication on ice and clarified by centrifugation at 21,000 x *g* for 30 min at 4°C. Cell lysate was filtered with 0.45 µM PES filters and incubated with Ni-NTA beads, equilibrated in J0 buffer supplemented with 500 mM NaCl, for 2 h at 4°C. Beads were sequentially washed in J0/500 mM NaCl, J0/750 mM NaCl/40 mM imidazole and in J0 buffer alone. RPA protein was eluted with J0 buffer supplemented with 500 mM imidazole. Eluate was dialysed into J0 buffer supplemented with 1 mM DTT overnight at 4°C. Dialysate was injected onto a Heparin HP 5 ml column (Cytiva) equilibrated in J0/1 mM DTT. RPA protein was eluted with a linear gradient from 0-1 M NaCl in J0/1 mM DTT and fractions containing RPA protein were pooled, concentrated, and further purified by SEC on a Superdex 200 column in 20 mM HEPES pH 7.5, 1 M NaCl, 1 mM DTT. Pure fractions (>90% by Coomassie Brilliant Blue-stained 10% SDS-PAGE and with an A280/A260 <0.6 to minimise DNA contamination) were pooled, concentrated and NaCl was diluted to ∼300 mM NaCl and glycerol added to ∼7.5% (v/v) before storage at −20°C (short-term) or −80°C (long-term). RPA complex point and truncated mutants were prepared in an equivalent manner. RPA was purified from bacteria >4 times and all protein preparations produced equivalent yields and activity in experimental assays.

GST-tagged versions of RPA70 and RPA32 domains were expressed from a pOPINK vector background. Transformed *E. coli* Rosetta2 (DE3)/pLysS were grown to OD_600_ 0.6 in LB broth and induced with 0.5 mM IPTG for 3 h at 37°C. Cells were harvested by centrifugation and re-suspended in lysis buffer (20 mM Tris pH 7.5, 500 mM NaCl, 0.1% Triton X-100 1x EDTA-free protease inhibitors (Roche)) and stored at −20°C until use. Cells were lysed by sonication on ice and clarified by centrifugation at 21,000 x *g* for 30 min at 4°C. Cell lysate was filtered with 0.45 µM PES filters and incubated with Glutathione Sepharose 4B beads, equilibrated in lysis buffer minus the protease inhibitors, for 2 h at 4°C. Beads were washed with wash buffer (20 mM Tris pH 7.5, 500 mM NaCl) and eluted in wash buffer supplemented with 20 mM reduced glutathione (Sigma) at pH 7.5. The eluate was concentrated and further purified by SEC using a Superdex 200 column and eluted by isocratic elution in 20 mM Tris pH 7.5, 100 mM NaCl, 1 mM DTT. Pure fractions (>90% by Coomassie Brilliant Blue-stained 10% SDS-PAGE) were pooled, concentrated and glycerol was added to 10% (v/v) before storage at −20°C (short-term) or −80°C (long-term).

Human PCNA was cloned into and expressed from a pOPINB vector background with an N-terminal His-3C tag. Transformed *E. coli* Rosetta2 (DE3)/pLysS were grown to OD_600_ 0.6 in LB broth and induced with 0.5 mM IPTG for 3 h at 37°C. Cells were harvested by centrifugation and re-suspended in lysis buffer (20 mM Tris pH 7.5, 500 mM NaCl, 20 mM imidazole, 0.1% Triton X-100 1x EDTA-free protease inhibitors (Roche)) and stored at −20°C until use. Cells were lysed by sonication on ice and clarified by centrifugation at 21,000 x *g* for 30 min at 4°C. Cell lysate was filtered with 0.45 µM PES filters and incubated with Ni-NTA beads (Thermo Scientific), equilibrated in lysis buffer minus the protease inhibitors, for 2 h at 4°C. Beads were sequentially washed with high salt (20 mM Tris pH 7.5, 500 mM NaCl, 20 mM imidazole) and low salt wash buffer (20 mM Tris pH 7.5, 100 mM NaCl, 20 mM imidazole) before re-suspension in low salt wash buffer. His-3C (made in-house) was added to the re-suspended beads and incubated overnight at 4°C with rotation. The subsequent flow-through containing untagged proteins (with an additional N-terminal Gly-Pro following 3C digestion) was further purified by SEC in 20 mM Tris pH 7.5, 100 mM NaCl, 1 mM DTT. Pure fractions (>90% by Coomassie Brilliant Blue-stained 10% SDS-PAGE) were pooled, concentrated and glycerol was added to 10% (v/v) before storage at −20°C (short-term) or −80°C (long-term).

Ub chains made in-house were essentially prepared as described ^56^. Briefly, E1-activating enzyme (mE1) was expressed in *E. coli* from a pET vector with an N-terminal His_6_-tag before purification using Ni-NTA beads and anion-exchange chromatography. E2-conjugating enzymes (Ubc13, Uev1a and Cdc34) were expressed in *E. coli* from pGEX6P-1 vectors with an N-terminal GST-3C tag before purification using Glutathione Sepharose 4B beads and elution by cleaving the tag with GST-tagged 3C protease on the beads. Ub was expressed in *E. coli* from a pET17b vector without a tag. After cell lysis as above, perchloric acid was slowly added to 0.5% (v/v) with stirring on ice and the supernatant after centrifugation was dialysed into 50 mM sodium acetate pH 4.7. The dialysate was injected onto 2 x 5 ml HiTrap SP FF columns (Cytiva) connected in tandem and eluted with a linear gradient from 0-1 M NaCl in 50 mM sodium acetate pH 4.7. Peak fractions were analysed by 17.5% SDS-PAGE and pure protein was pooled and dialysed into 10 mM Tris pH 7.5. PolyUb chains were prepared enzymatically as described ^56^ and purified using a Resource S 6 ml column (Cytiva). For preparing branched chains, K63-linked tri- and tetra-Ub chains were prepared using Ubc13/Uev1a with *Δ*GlyGly and K48R Ub mutants followed by purification using a Resource S 6 ml column. Subsequent formation of a branched chain was carried out using Cdc34 and K48R Ub and similar purification step to remove Ub monomers.

### GST pull-downs

2.5 or 5 µg GST or GST-tagged proteins were immobilised to either Glutathione Sepharose 4B beads or glutathione magnetic agarose beads (Pierce) in binding/wash buffer (20 mM Tris pH 7.5, 150 mM NaCl, 0.01% IGEPAL CA-630, 1 mM DTT) before incubation with 10 µg untagged RPA or ZUP1 protein was added in a total volume of 400 µl binding buffer for 1 h at 4°C with rotation. Beads were washed twice with 400 µl binding/wash buffer and proteins were eluted by boiling in 20 µl SDS loading buffer (50 mM Tris pH 6.5, 2% SDS, 10% glycerol, 50 mM DTT, bromophenol blue). Pulldowns were analysed by SDS-PAGE with 5-20% input (indicated for each experiment) and 50% of the pulldown loaded per gel followed by Coomassie Brilliant Blue staining or immunoblotting against the indicated antibodies. Identical results were obtained with both bead types used.

For quantification of the GST pulldowns with RPA fragments, bands corresponding to RPA70 were quantified using ImageLab (Bio-Rad) and normalised to the band intensity of GST-ZUP1 in the corresponding lane. The binding of each RPA sample was quantified in this way in three independent experiments and normalised to the binding of wildtype RPA within the same experiment and gel. Bar charts of binding were prepared using GraphPad Prism 9 with the mean plotted and error bars representing the standard deviation.

### Electrophoretic mobility shift assay (EMSA)

0.25 µM ssDNA-containing substrate (Gap-26, formed using ctatggcgaggcgattatcaacccatttagtcgtaatagtgaagagtcacgacaacatcg, annealed with taatcgcctcgccatag and cgatgttgtcgtgactc) ^57^ was incubated with the indicated amounts of RPA complex and ZUP1 proteins in 20 mM Tris pH 7.5, 100 mM NaCl, 0.5 mg/ml BSA, 1 mM DTT in a volume of 10 µl. Samples were incubated for 5 min at 25°C and analysed by 5% native PAGE in 0.2x TBE running buffer at 4°C. Bands of free and shifted Gap-26 substrate were visualised with SYBR safe.

#### *In vitro* deubiquitination (DUB) assays

ZUP1 with GST or RPA, and poly-Ub chains (made in-house or from Bio-Techne or UbiQ) were incubated separately in DUB buffer (20 mM Tris pH 7.5, 100 mM NaCl, 10 mM DTT) for 5 min at 25°C before combining to give final concentration of 1 µM ZUP1, 1.25 µM GST or RPA (altered concentrations indicated in figure legends) and 2 µM poly-Ub chains or 1.75 µg longer (Ub6+) chains. End-point assays were incubated for 2 h at 25°C unless specified or as indicated in time course assays. Reactions were quenched with SDS-loading buffer without boiling and analysed by SDS-PAGE and Coomassie Brilliant Blue staining.

For semi-quantitative DUB assays, a time course was set up as above and samples were taken at 5, 15, 30, 60, and 120 min intervals and analysed by SDS-PAGE and Coomassie Brilliant Blue staining overnight. Gels were scanned with the Gel Doc XR+ (BioRad). Bands corresponding to tetra-Ub were quantified using ImageLab (Bio-Rad) relative to the tetra-Ub input band for each independent assay within the same gel. The proportion of tetra-Ub cleaved in the assay was calculated by subtracting the relative amount of tetra-Ub present from 1. This was carried out in three independent experiments for each ZUP1 protein -/+ RPA and plotted using GraphPad Prism 9 with error bars representing the standard deviation. A non-linear regression curve was fit with one phase exponential decay.

For quantitative DUB assays with a Ub-AMC minimal substrate, master mixes of ZUP1, RPA or GST and Ub-AMC were prepared before applying to 96-well half area flat bottom black plates (Corning). ZUP1 was at a final constant concentration of 1 µM and GST or RPA at 1.25 µM (except in titration assays) before adding Ub-AMC at the indicated final concentrations just before starting the fluorescence reading. Measurements were taken every 1 min for 60 min at 37°C with an excitation wavelength of 360 nm and emission wavelength of 460 nm using an LB941 TriStar plate reader. AMC (Bio-Techne) standards were prepared by serial dilution in DUB buffer. Measurements for the enzyme kinetics and for the RPA titration with WT ZUP1 were carried out three times. The fluorescence output (RFU) was converted to [AMC] released using the AMC standard curve after normalising to buffer-only blank wells. [AMC] released was plotted against time for each Ub-AMC concentration for each ZUP1 protein with GST or RPA. The [AMC] released over the starting 4 min could be fitted with a linear fit and was used to calculate V_0_ (initial rate) for ZUP1 DUB activity against the minimal substrate, which was plotted against Ub-AMC concentration with error bars representing the standard error of the mean (SEM) for each point. As the resulting V_0_ v Ub-AMC concentration data could not be fitted with Michaelis-Menten kinetics, a linear fit model could be applied and the gradient of this was used to compare ZUP1 DUB activity with GST or RPA as the Specific Rate (nM/min/µM), which was plotted as a bar graph with the error bars representing how well the linear model fits the data. For the RPA titration experiments, 1 µM ZUP1, 1 µM Ub-AMC and a titration of RPA was used in three separate experiments and the initial rate (V_0_) was plotted against RPA concentration with error bars representing the SEM. A one phase exponential decay non-linear regression model was applied to provide an apparent k_d_ for RPA binding to stimulate ZUP1 DUB activity.

In assays using Ub-propargylamine (Ub-PA, UbiQ), USP2 (Bio-Techne) and ZUP1 proteins were pre-incubated in DUB buffer as above separately from Ub-PA for 5 min at 25°C before mixing to give final concentration of 1 µM USP2/GST/ZUP1 and 3 µM Ub-PA. Reactions were incubated for 3 h at 25°C and quenched with SDS-loading buffer without boiling. Reactions were analysed by SDS-PAGE and Coomassie Brilliant Blue staining.

### Sample preparation for XL-MS

Samples were buffer matched into 20 mM HEPES-KOH pH 7.5, 100 mM NaCl, 1 mM DTT by dialysis and incubated after mixing for 15 min at 25°C. 30 µg of ZUP1 and 50 µg of RPA (ZUP1 in slight molar excess) were used in each experiment with a final protein concentration of approximately 1 mg/ml. For experiments with K63-linked Ub4 chains, 20 µg was added so as to be in a slight molar excess to ZUP1. A 1 mg aliquot of isotopically coded BS3 d_0_/d_12_ crosslinker (Creative Molecules, 001SS) was reconstituted to 25 mM in water and immediately added to the mixture in equimolar amounts to the number of moles of lysine residues present in the sample. The crosslinking reaction was incubated for 30 min at 25°C with mild agitation and the reaction was quenched with 50 mM NH_4_HCO_3_ for 10 min at 25°C. The samples were reduced in volume in a speedvac and re-suspended in 50 µl 7.2 M deionised urea, 100 mM NH_4_HCO_3_ and incubated for 15 min at 25°C. Cysteines were reduced with 10 mM DTT for 1 h at 25°C and cysteine residues were protected with 50 mM iodoacetamide (Sigma, I1149-25G) for 45 min in the dark at 25°C. The reaction was quenched with DTT and urea was diluted to <1 M with 50 mM NH_4_HCO_3_ before adding MS-grade trypsin (Promega, V5113) in a ratio of 1:25 (trypsin:protein) and incubated for 18 h at 37°C. The reaction was stopped by the addition of formic acid (0.5% final concentration, v/v) and peptides were purified using C18 stage tips (Glygen). Peptides were eluted in 60% acetonitrile, 0.1% formic acid and dried and stored at −20°C.

For experiments using strong cation exchange (SCX) chromatography to enrich for highly charged crosslinked peptide species, dried peptides following desalting were re-suspended in 20% acetonitrile, 0.4% formic acid and purified using initialised SCX Stage Tips (Thermo-Scientific). After washing in 20% acetonitrile, 0.4% formic acid, crosslinked peptides were eluted by stepped elution in 15 mM, 20 mM, 40 mM, 70 mM, 125 mM and 500 mM ammonium acetate (all in 20% acetonitrile, 0.4% formic acid). Eluates were dried and stored at −20°C.

### LC-MS/MS and analysis

Peptides were separated by nano liquid chromatography (Thermo Scientific Ultimate RSLC 3000) coupled in line a Q Exactive mass spectrometer equipped with an Easy-Spray source (Thermo Fischer Scientific). Peptides were trapped onto a C18 PepMac100 precolumn (300 µm i.d.x5mm, 100 Å, ThermoFischer Scientific) using Solvent A (0.1% Formic acid, HPLC grade water). The peptides were further separated onto an Easy-Spray RSLC C18 column (75 µm i.d., 50 cm length, Thermo Fischer Scientific) using a 120 min linear gradient (15% to 35% solvent B (0.1% formic acid in acetonitrile)) at a flow rate 200 nl/min. The raw data were acquired on the mass spectrometer in a data-dependent acquisition mode (DDA). Full-scan MS spectra were acquired in the Orbitrap (Scan range 350-1500 m/z, resolution 70,000; AGC target, 3e6, maximum injection time, 50 ms). The 10 most intense peaks were selected for higher-energy collision dissociation (HCD) fragmentation at 30% of normalized collision energy. HCD spectra were acquired in the Orbitrap at resolution 17,500, AGC target 5e4, maximum injection time 120ms with fixed mass at 180 m/z. Charge exclusion was selected for unassigned and 1+ ions. The dynamic exclusion was set to 40 s.

Raw files were analysed directly using pLink2.0 ^58^ to identify crosslinked peptides. BS3 and BS3-d12 (manually input) were used as the designated crosslinkers. Cys carbamidomethylation was used as a fixed modification and Met oxidation and Gln/Asn deamidation as variable modifications. Search parameters included peptide length of 4-100 and peptide mass of 400-10,000, maximum 3 missed trypsin cleavages allowed and a precursor and fragment mass tolerance of 20 ppm. An FDR of < 1% was used to filter out low-confidence crosslinks. Fasta files containing the amino acid sequences of the relevant proteins were used alongside a contaminant *E. coli* database. The resulting high-confidence crosslinks were further manually filtered by only including crosslinks with evidence in multiple samples in at least one of the technical replicates. The remaining crosslinks were visualised using xVis ^59^. Crosslinks were superimposed onto the previously published ZUP1 structure (PDB: 6FGE) using ChimeraX ^60^. Excel tables of the crosslinked sites identified can be found in Supplementary Data File 1. XL-MS data (raw files and raw output from pLink2.0) have been uploaded to ProteomeXchange via the PRIDE repository with identifier PXD039482.

### Size-exclusion chromatography and SEC-MALS

Size-exclusion chromatography (SEC) experiments were carried out on an AKTA Pure system (Cytiva) using a Superdex 200 10/300 or Superose 6 10/300 column (Cytiva) that was calibrated using molecular weight markers (Cytiva). The proteins were mixed and incubated for 5 min on ice and clarified by centrifugation at maximum speed in a bench top centrifuge at 4°C, before loading onto the column. Protein and protein complexes were eluted using isocratic elution in 20 mM Tris-HCl pH 7.5, 100 mM NaCl, 1 mM DTT at 0.5 ml/min. For SEC-MALS, ZUP1 was dialysed into 20 mM Tris pH 7.5, 100 mM NaCl, 1 mM DTT and a serial dilution was loaded onto a Superdex 200 10/300 column at 22°C using a Shimadzu chromatography system connected in-line to a Heleo8+ multi angle light scattering detector and an Optilab T-rEX refractive index (RI) detector. Results were processed and analysed using ASTRA 6 (Wyatt Technologies).

## Supplementary items

**Supplementary data file 1.** Folder containing data tables of the crosslinks identified using XL-MS.

**Supplementary data file 2.** Folder containing examples of spectra derived from XL-MS experiments.

## Notes

### Competing Interest Statement

The authors have declared no competing interest.

