## Extended Data Figures for "The RPA complex orchestrates K63-linked deubiquitination via ZUP1"

### Extended Data Fig 1

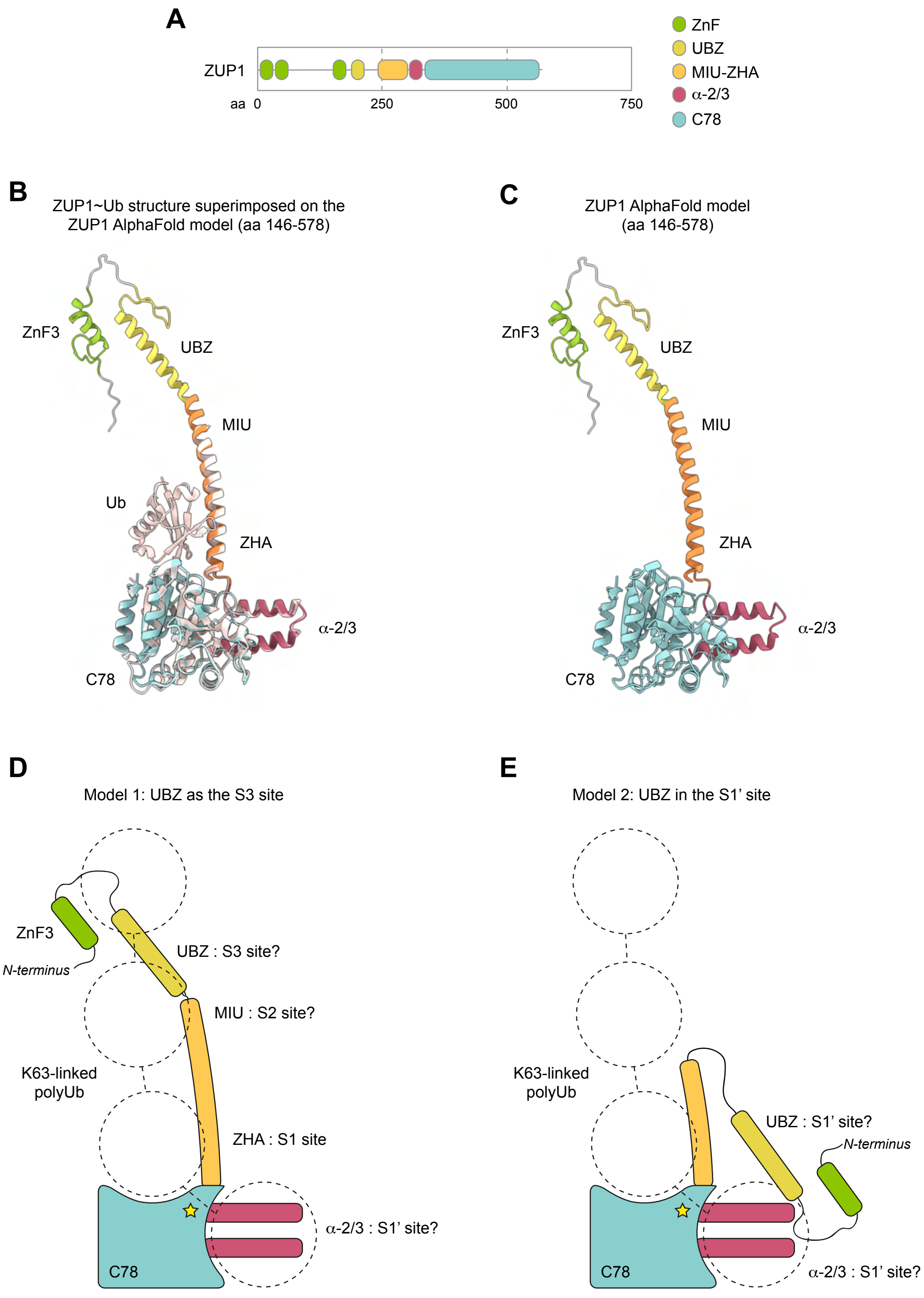

#### Extended Data Fig 1

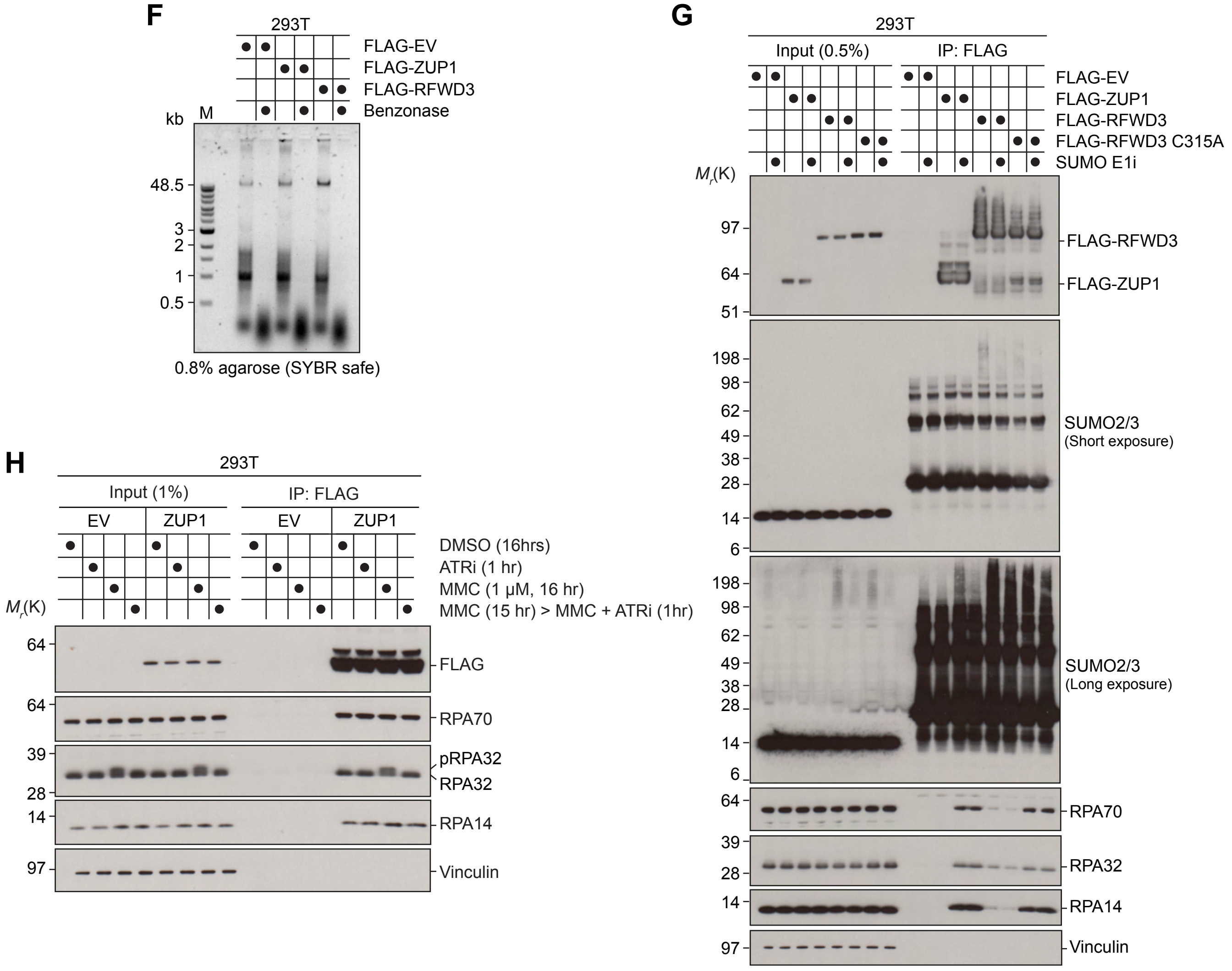

**Extended Data Figure 1. ZUP1 and RPA form a complex in human cells.**

**A.** Domain organisation of human ZUP1.

**B.** Structural superposition of ZUP1~Ub (light pink, PDB: 6FGE) on to the AlphaFold model of ZUP1 (AF-Q96AP4-F1), with the domains coloured as in (A). Note that only aa 146-578 of the AlphaFold ZUP1 model are shown for clarity.

**C.** AlphaFold model of ZUP1 146-578 only.

**D.** Model depicting how the ZUP1 UBZ domain might act as the S3 site for a K63-linked polyUb chain.

**E.** Model depicting how the ZUP1 UBZ domain might contribute to the S1' site for a K63-linked polyUb chain.

**F.** Cells were lysed in the absence (-) or presence (+) on benzonase (>500 units/ml). DNA was extracted from the lysates and analysed by DNA gel electrophoresis to verify the benzonase activity.

**G.** 293T cells were transfected with FLAG-EV, FLAG-ZUP1, FLAG-RFWD3, or FLAG-RFWD3 C315A and then treated with 1 μM SUMO E1i for 90 mins. Cells were lysed, subjected to FLAG immunoprecipitation (IP), and analysed by immunoblotting with the indicated antibodies. Data is representative from $n = 2$ biological repeats.

**H.** 293T cells were transfected with FLAG-EV or FLAG-ZUP1 and then treated with 1 μM MMC for 16 h, 5 μM ATRi for 1 h, or MMC/ATRi and then processed as in (B).

#### Extended Data Fig 2

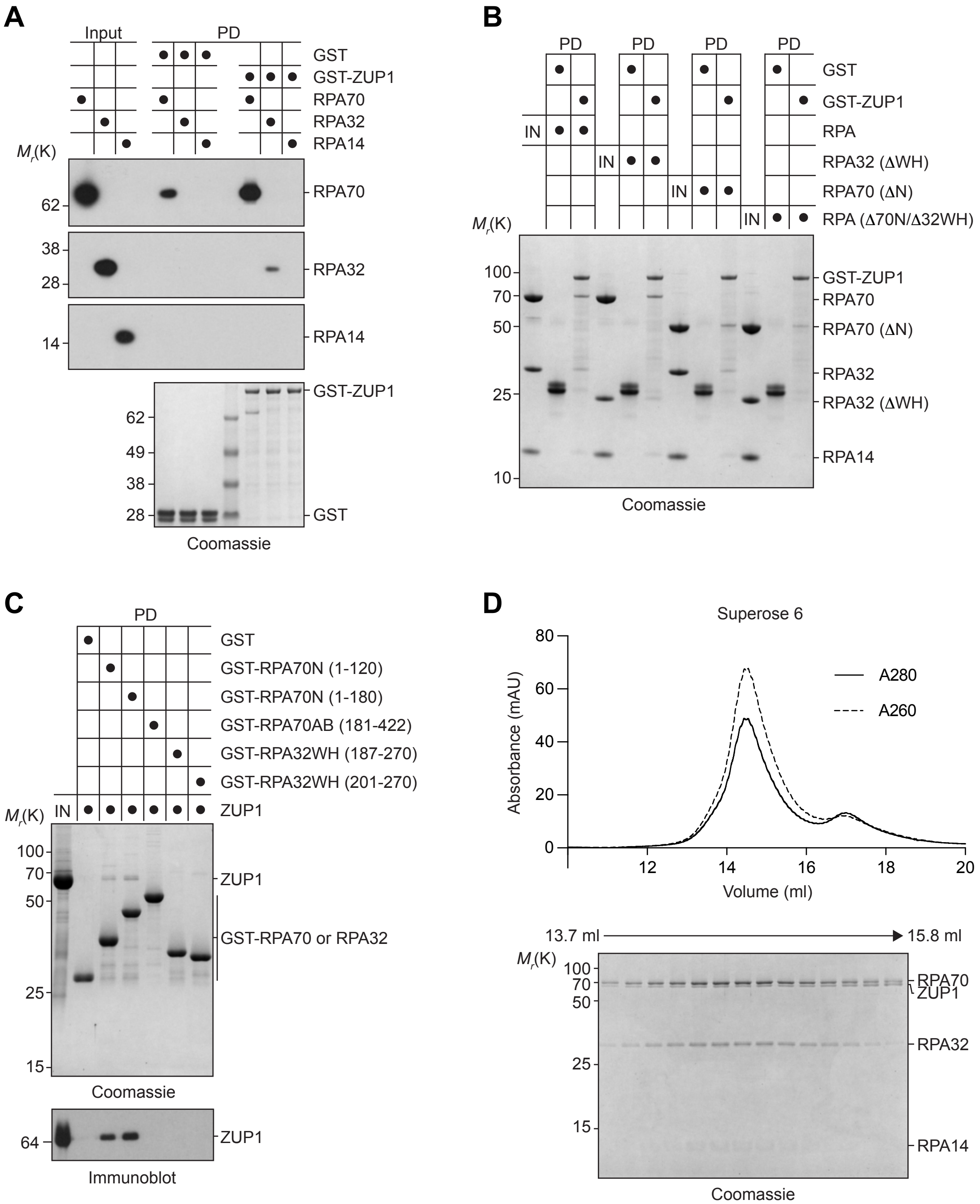

**Extended Data Figure 2. ZUP1 and RPA form a complex *in vitro*.**

- **A.** GST pulldown (PD) assay using GST and GST-ZUP1 incubated with *E. coli* cell lysate containing recombinantly expressed untagged RPA70, RPA32, or RPA14. Inputs and PDs were analysed by SDS-PAGE followed by Coomassie staining or immunoblotting with the indicated antibodies.
- **B.** GST PD assay using GST or GST-tagged ZUP1 with either the wild type RPA complex or the indicated truncated RPA complexes. 10% of the input and 50% of the PD was analysed by SDS-PAGE followed by Coomassie staining or immunoblotting with ZUP1 antibody.
- **C.** GST PD assay with GST or GST-RPA70 or GST-RPA32 fragments. 10% of the input and 50% of the PD was analysed by SDS-PAGE followed by Coomassie staining.
- **D.** Chromatogram showing the absorbance at 280 and 260 nm of RPA, ssDNA (dt100) and ZUP1 run over a Superose 6 10/300 gel filtration column. Proteins were eluted and peak fractions were separated by SDS-PAGE gel and analysed by Coomassie staining. The absorbance at 260 nm indicates the presence of ssDNA within the complex. 100 µg of each RPA and ZUP1 were mixed with 6.1 µg dt100 leading to a molar ratio of 1:1.7:0.25, RPA:ZUP1:dt100, resulting in a molar excess of ZUP1 over RPA-ssDNA complex. Free ZUP1 can be detected in the second, smaller peak indicating approximately 1:1 (ZUP1:RPA-ssDNA) stoichiometry in the main peak.

**Extended Data Fig 3**

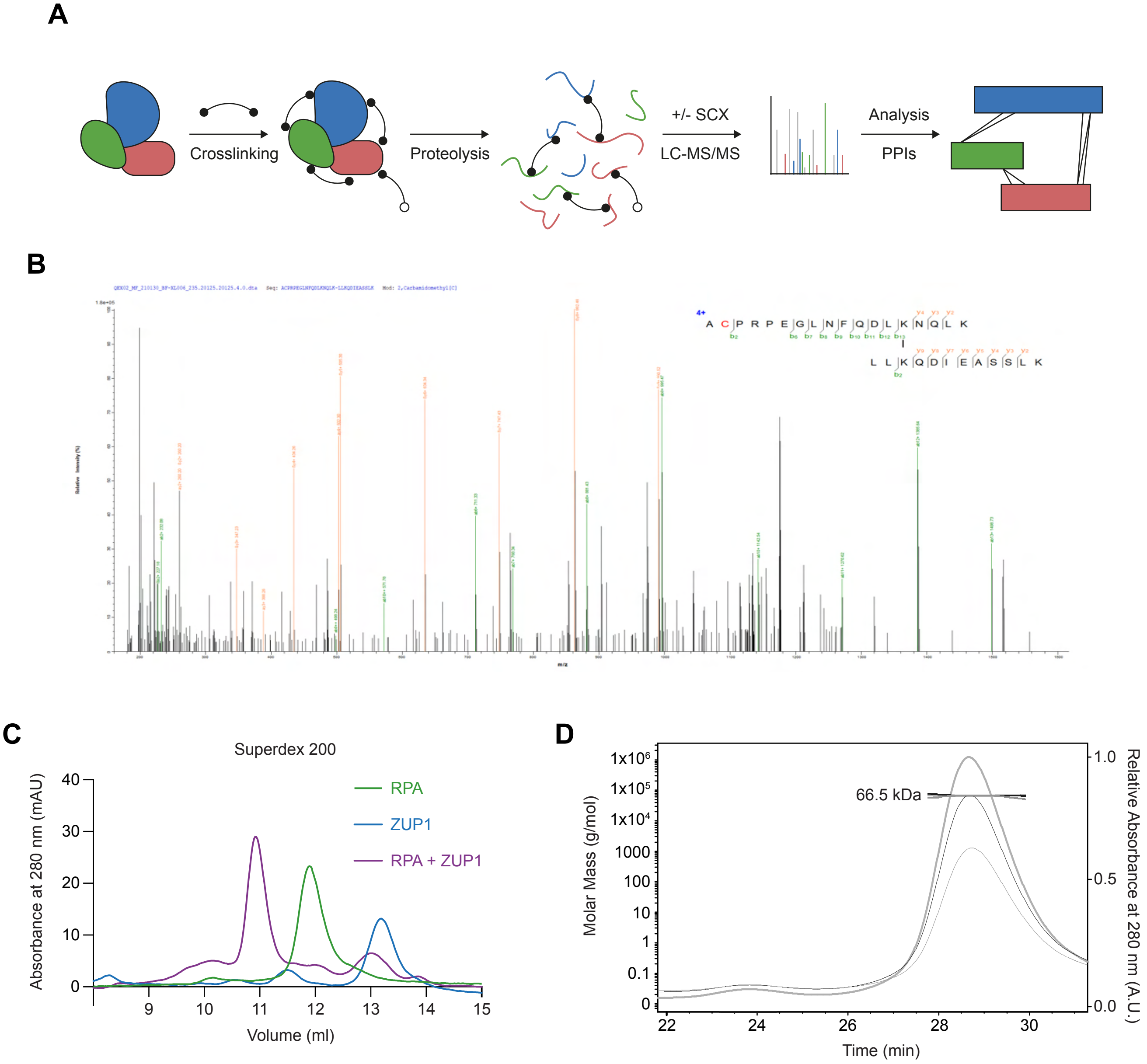

**Extended Data Figure 3. XL-MS analysis of the ZUP1-RPA complex.**

- **A.** XL-MS workflow used in this study.
- **B.** Example spectra from the XL-MS workflow. Full tables of crosslinks and the output from pLink2.0 can be found in Supplemental Data File 1 with further example spectra in Supplemental Data File 2
- **C.** Chromatogram showing the absorbance at 280 nm of RPA or ZUP1 alone or RPA and ZUP1 run over a Superdex 200 10/300 gel filtration column. 200 µg of each were mixed and loaded, resulting in a ~1.7 molar excess of ZUP1 over RPA. Free ZUP1 can be detected in the second, smaller peak in the complex indicating approximately 1:1 stoichiometry.
- **D.** SEC-MALS of ZUP1 indicating that is it a monomer at ~66 kDa in solution at the concentrations used in the binding, DUB, and XL-MS assays in this study.

#### Extended Data Fig 4

A

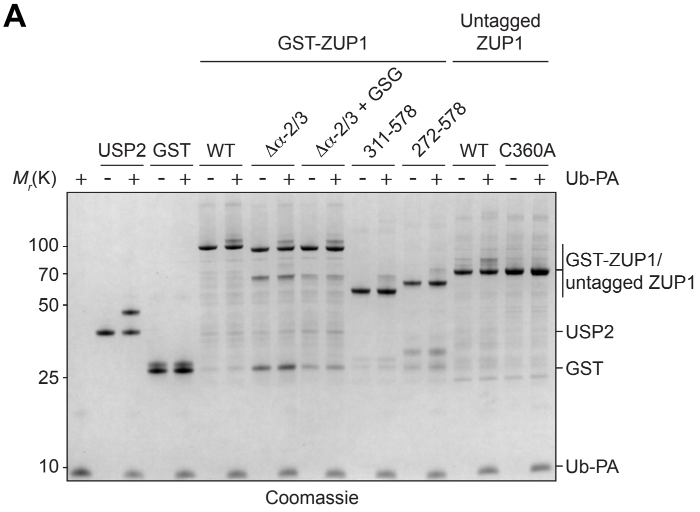


B

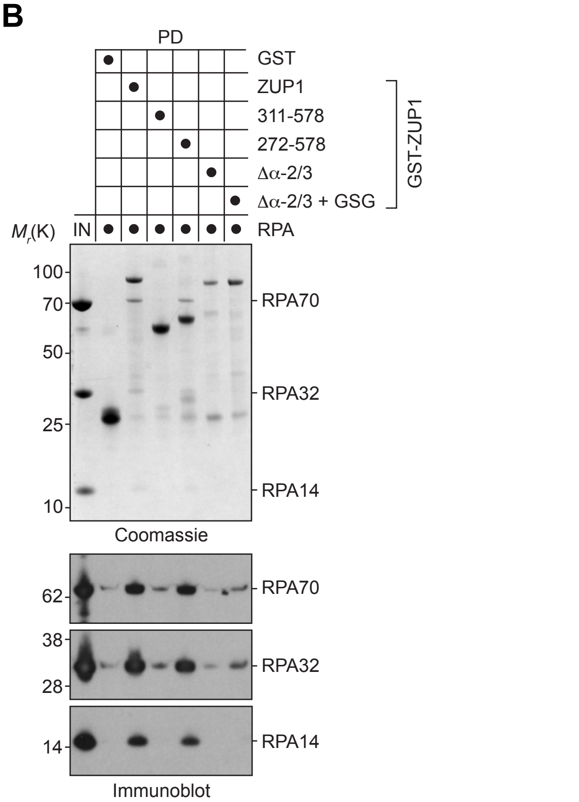


C

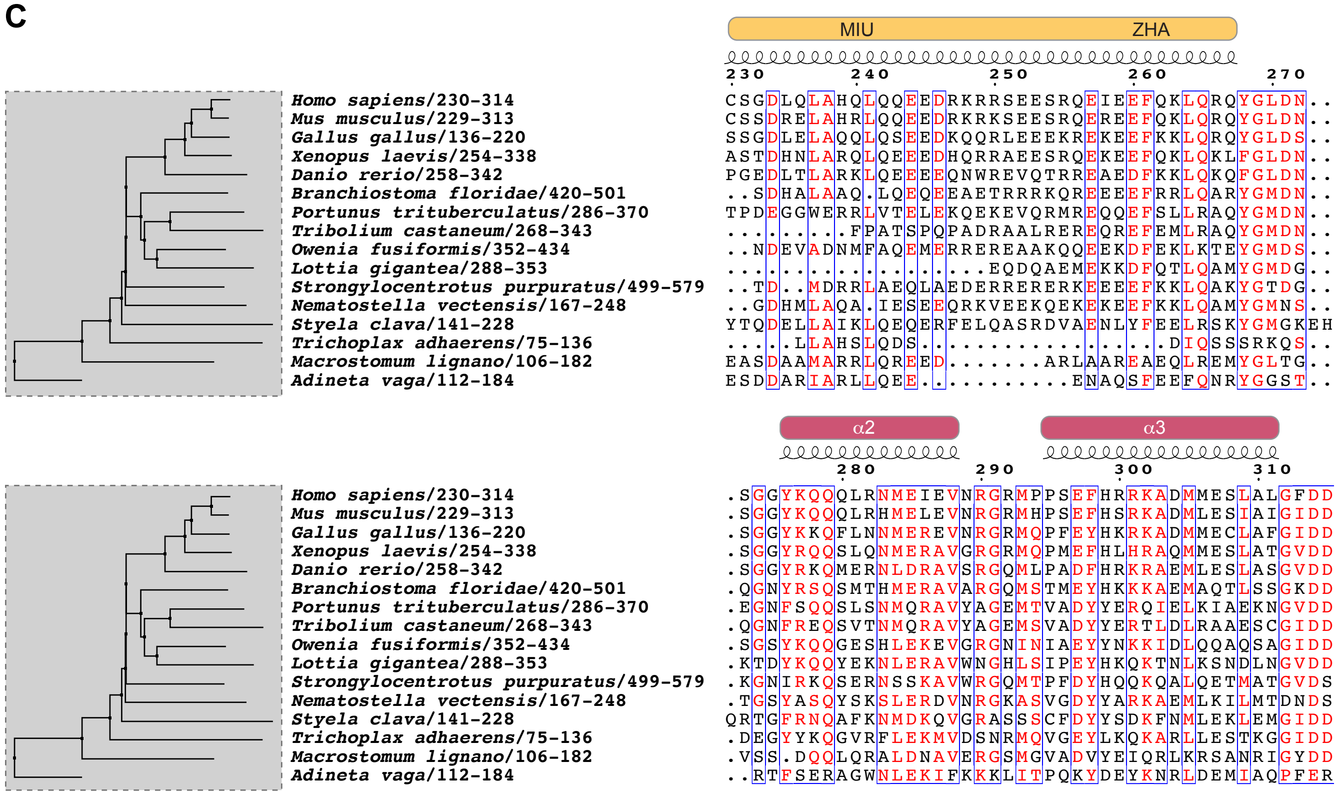

#### Extended Data Fig 4

**D**

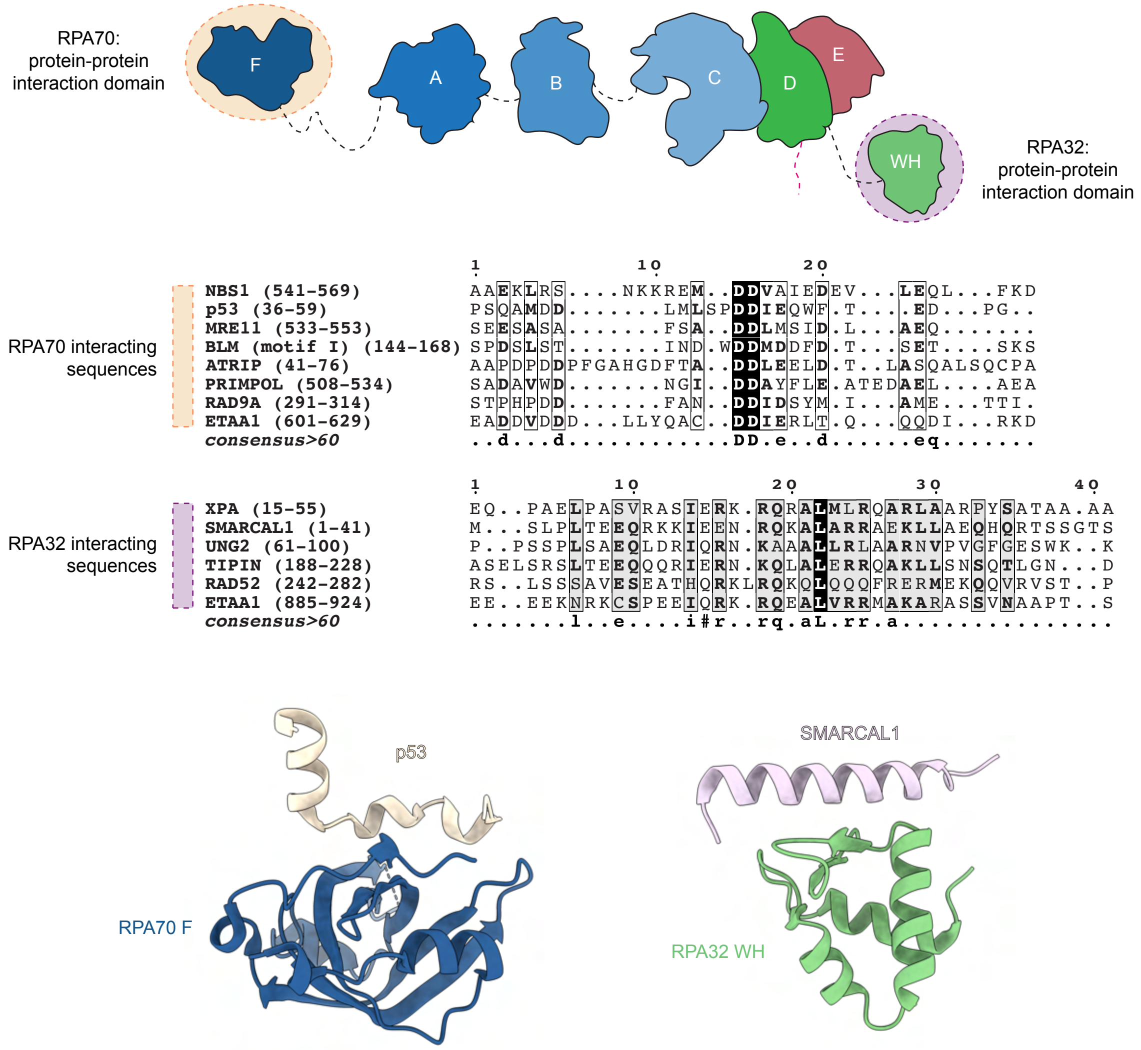

**Extended Data Figure 4. The ZUP1 α-2/3 region promotes the formation of the ZUP1-RPA complex.**

**A.** The indicated proteins were reacted with Ub-PA for 3 h at 25°C. Reaction products were analysed by SDS-PAGE followed by Coomassie staining. ZUP1 C360A did not react with Ub-PA as expected, but the remaining ZUP1 proteins show a band migrating slower than the unmodified protein indicating the catalytic cysteine is accessible for reacting. USP2 was used as a positive control.

**B.** GST pulldown (PD) assay with GST and GST-tagged ZUP1 proteins with untagged RPA. 10% of the input (IN) and 50% of the eluted PD were analysed by SDS-PAGE followed by Coomassie staining or immunoblotting with the indicated antibodies.

**C.** Multiple sequence alignment and evolutionary distances of the indicated ZUP1 homologs. Secondary structure and corresponding regions in *Hs*ZUP1 are shown.

**D.** *Top*, cartoon illustrating the domains within the RPA complex: RPA70 consists of domains F, A, B, and C; RPA32 consists of domains D and WH; RPA14 consists of domain E. The RPA70 N-terminal F domain and the RPA32 WH domains mediate protein-protein interactions with RPA client proteins. The RPA32 N-terminal region is shown in dashed pink, whilst flexible interdomain connecting residues are shown in dashed black. *Middle*, multiple sequence alignments of known RPA70 and RPA32 binding proteins, whose RPA-interacting sequences have been defined. *Bottom*, published structures of the RPA70 F domain with a p53 peptide (PDB: 2B3G) and the RPA32 WH domain with a SMARCAL1 peptide (PDB: 4MVQ).

**Extended Data Fig 5**

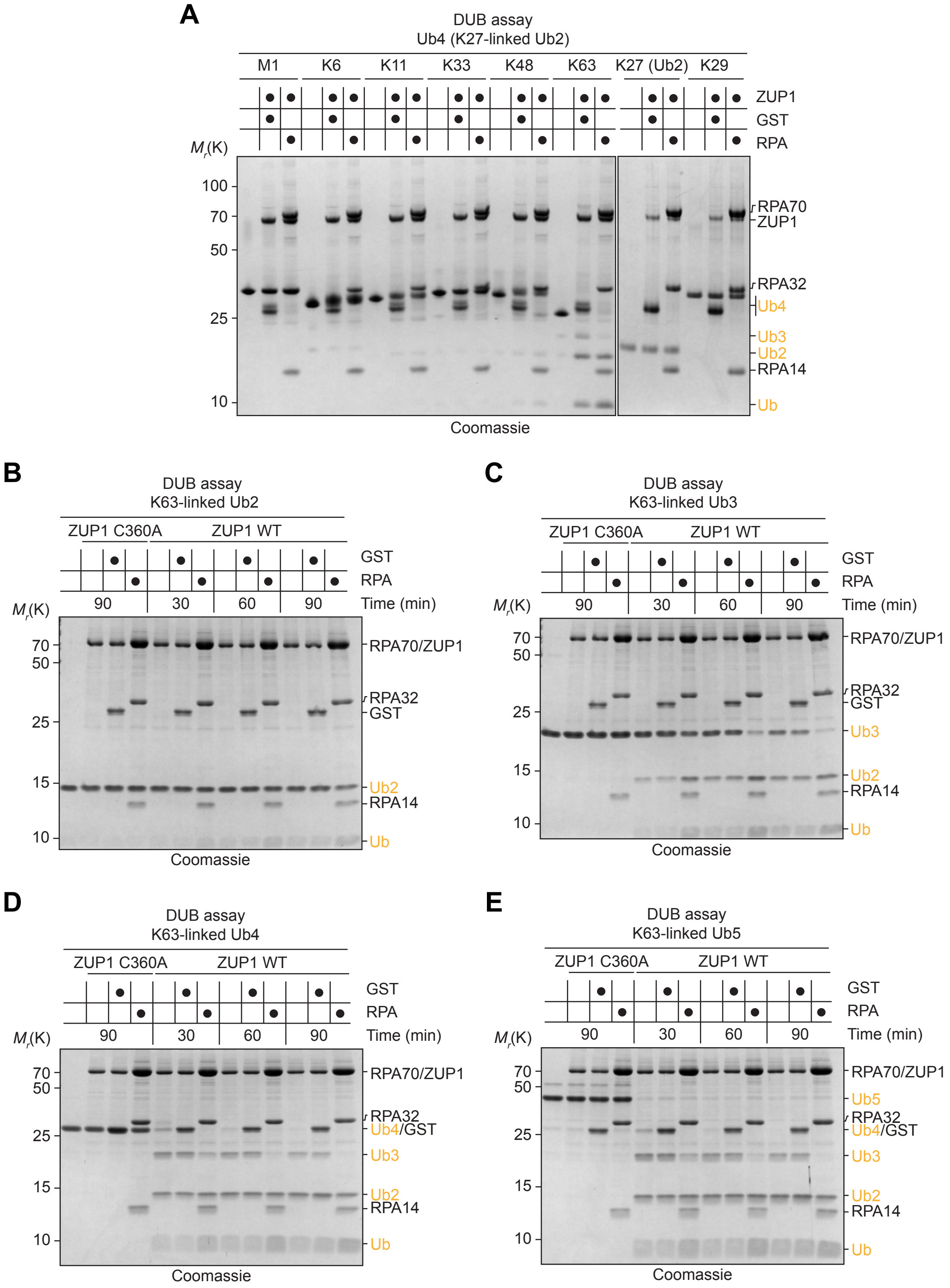

### Extended Data Fig 5

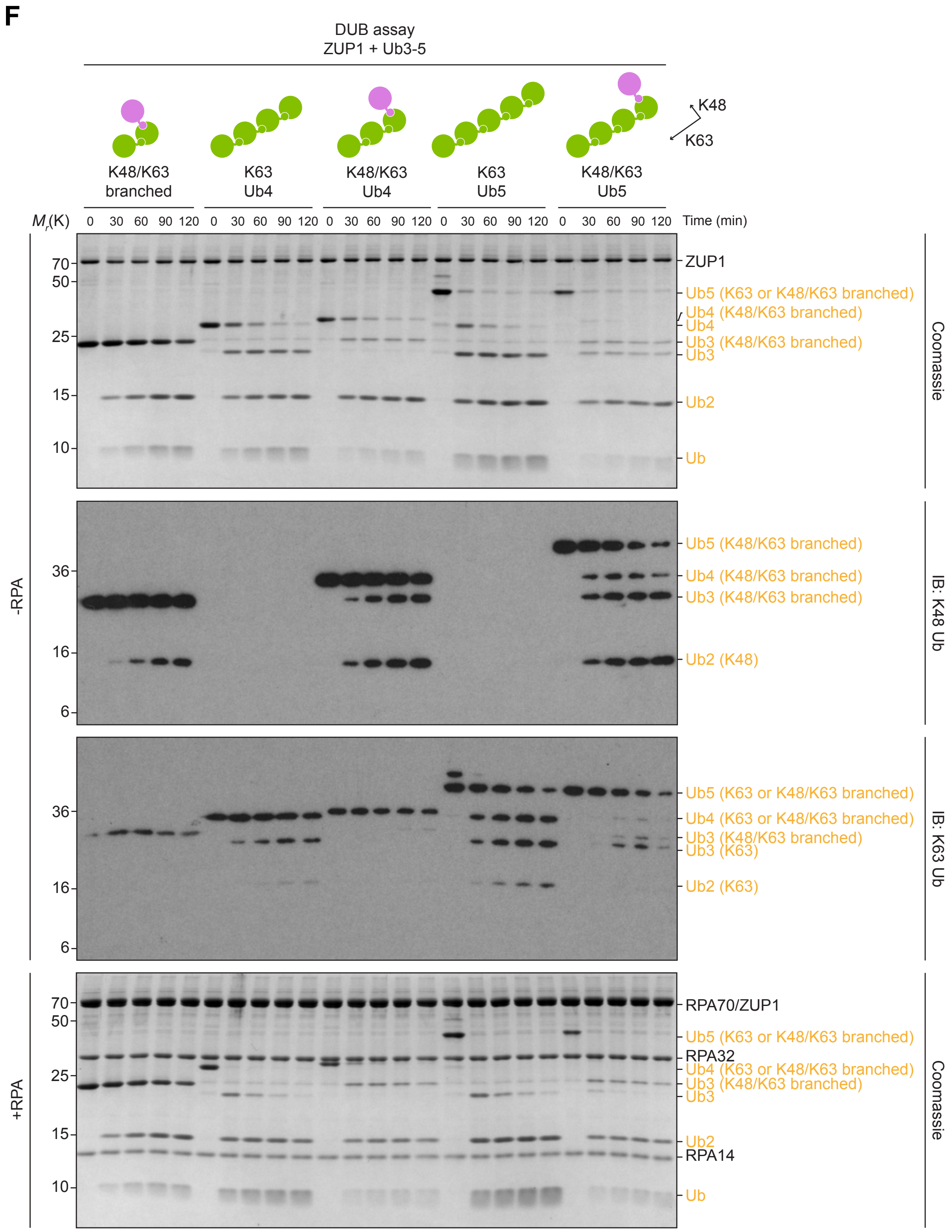

**Extended Data Fig 5**

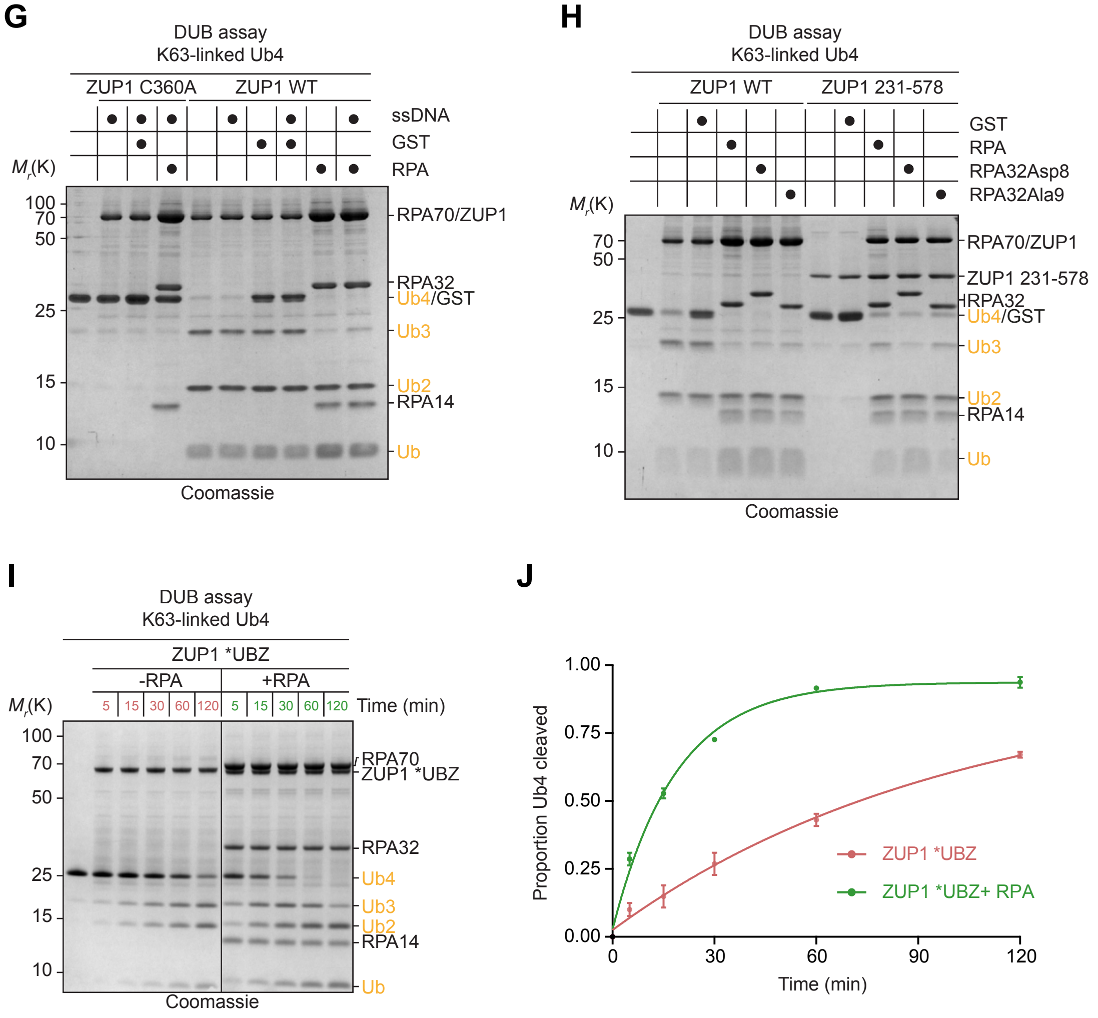

### Extended Data Fig 5

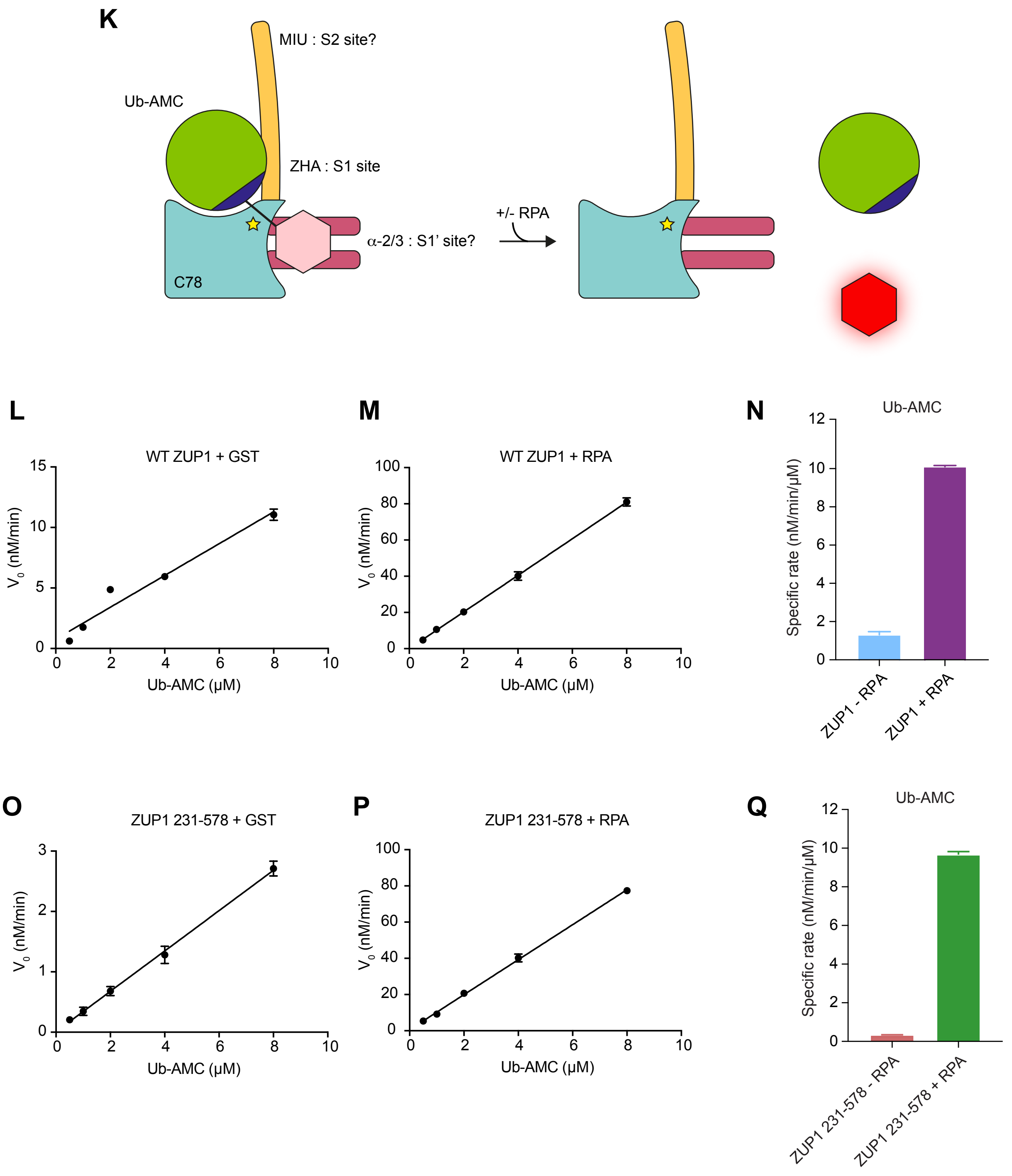

**Extended Data Figure 5. RPA stimulates ZUP1 K63-linked deubiquitination activity.**

**A.** *In vitro* DUB assay with the indicated following components: GST (1.25 μM), ZUP1 (1 μM), Ub4/Ub2 chains (2 μM), and RPA (1.25 μM). The reaction was incubated for 2 h at 25°C and quenched by the addition of SDS loading buffer without boiling. The reaction products were separated SDS-PAGE and stained with Coomassie.

**B.** As for (A), except with K63-linked Ub2 and inclusion of ZUP1 C360A.

**C.** As for (A), except with K63-linked Ub3 and inclusion of ZUP1 C360A.

**D.** As for (A), except with K63-linked Ub4 and inclusion of ZUP1 C360A.

**E.** As for (A), except with K63-linked Ub5 and inclusion of ZUP1 C360A.

**F.** *In vitro* DUB assay using homotypic K63-linked Ub chains or heterotypic K48/K63-linked Ub chains of differing lengths. Reaction products were analysed by SDS-PAGE followed by Coomassie staining or immunoblotting with the indicated antibodies.

**G.** As for (D), except with the inclusion ssDNA (0.25 μM). Different ssDNA-containing substrates gave similar results (data not shown).

**H.** As for (D), except with the indicated RPA complex mutants.

**I.** As for (D), except with ZUP1 *UBZ mutant containing I207A/L208A point mutants.

**J.** Quantification of the Ub4 band from (I). Data is from $n$ = 3 independent experiments and represents the mean and standard deviation.

**K.** Ub-AMC assay overview in the context of ZUP1 231-578. The Ub moiety in Ub-AMC interacts with the ZUP1 ZHA S1 site.

**L.** Ub-AMC was incubated with the indicated combinations of proteins at 37°C for 1 h with readings taken every 1 minute for AMC/fluorescence release. After a standard curve of AMC was used to normalise and convert the fluorescence output (RFU) of ZUP1 DUB activity against Ub-AMC, the initial rate ($V_0$) was calculated as the rate of AMC release per min for each Ub-AMC concentration over the initial 4 minutes of the reaction where the data could be modelled as being linear. The error bars are the standard error of the mean for each Ub-AMC concentration. The gradient of the initial rate linear fit was used to plot the Specific Rate. The ZUP1 C360A mutant was used as a negative control in these experiments.

**M.** As for (L), except with RPA.

**N.** The 'Specific Rate' is the initial rate for release of AMC from Ub-AMC per amount of Ub-AMC substrate provided. Error bars represent the closeness of the linear fit for the specific rate.

**O.** As for (L), except for ZUP1 231-578.

**P.** As for (M), except for ZUP1 231-578.

**Q.** As for (N), except for ZUP1 231-578.

#### Extended Data Fig 6

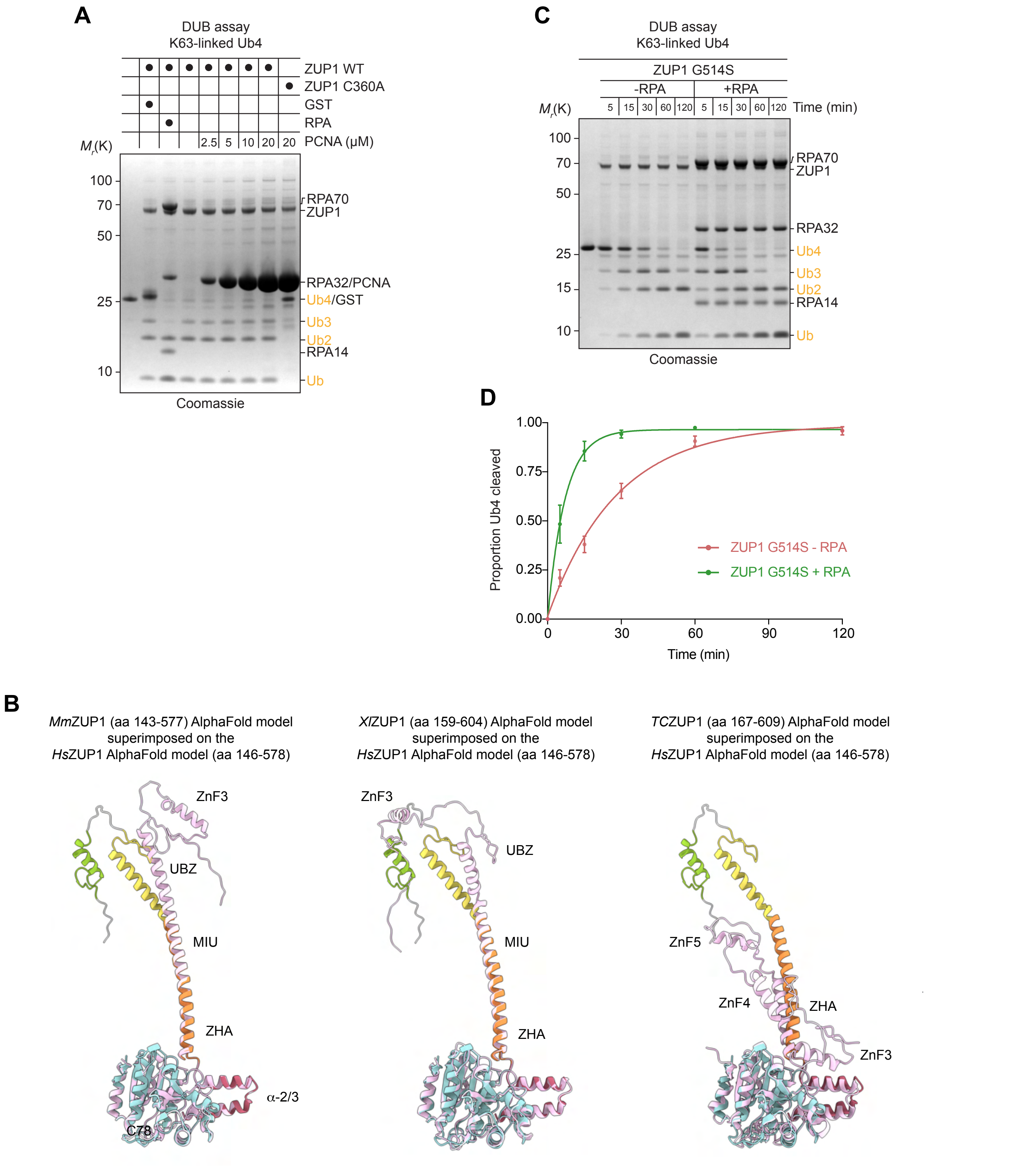

**Extended Data Figure 6. RPA is the main mechanism to stimulate ZUP1 deubiquitinase activity and is evolutionarily conserved.**

- **A.** *In vitro* DUB assay with the indicated following components: GST (1.25 µM), ZUP1 WT or C360A (1 µM), K63-linked Ub4 chains (2 µM), RPA (1.25 µM), and PCNA at the molarities shown in the figure. The reaction was incubated for 2 h at 25°C and quenched by the addition of SDS loading buffer without boiling. The reaction products were separated SDS-PAGE and stained with Coomassie.
- **B.** Structural superposition of AlphaFold models of ZUP1 orthologs on to the AlphaFold model of *Hs*ZUP1 aa 146-578. Domain names and locations are specific to the homolog.
- **C.** As in (A), except with the ZUP1 G514S mutant and samples were taken at the indicated times.
- **D.** Replotting of the G514S data presented in Figure 7C and D into one graph, instead of across two.

**Extended Data Fig 7**

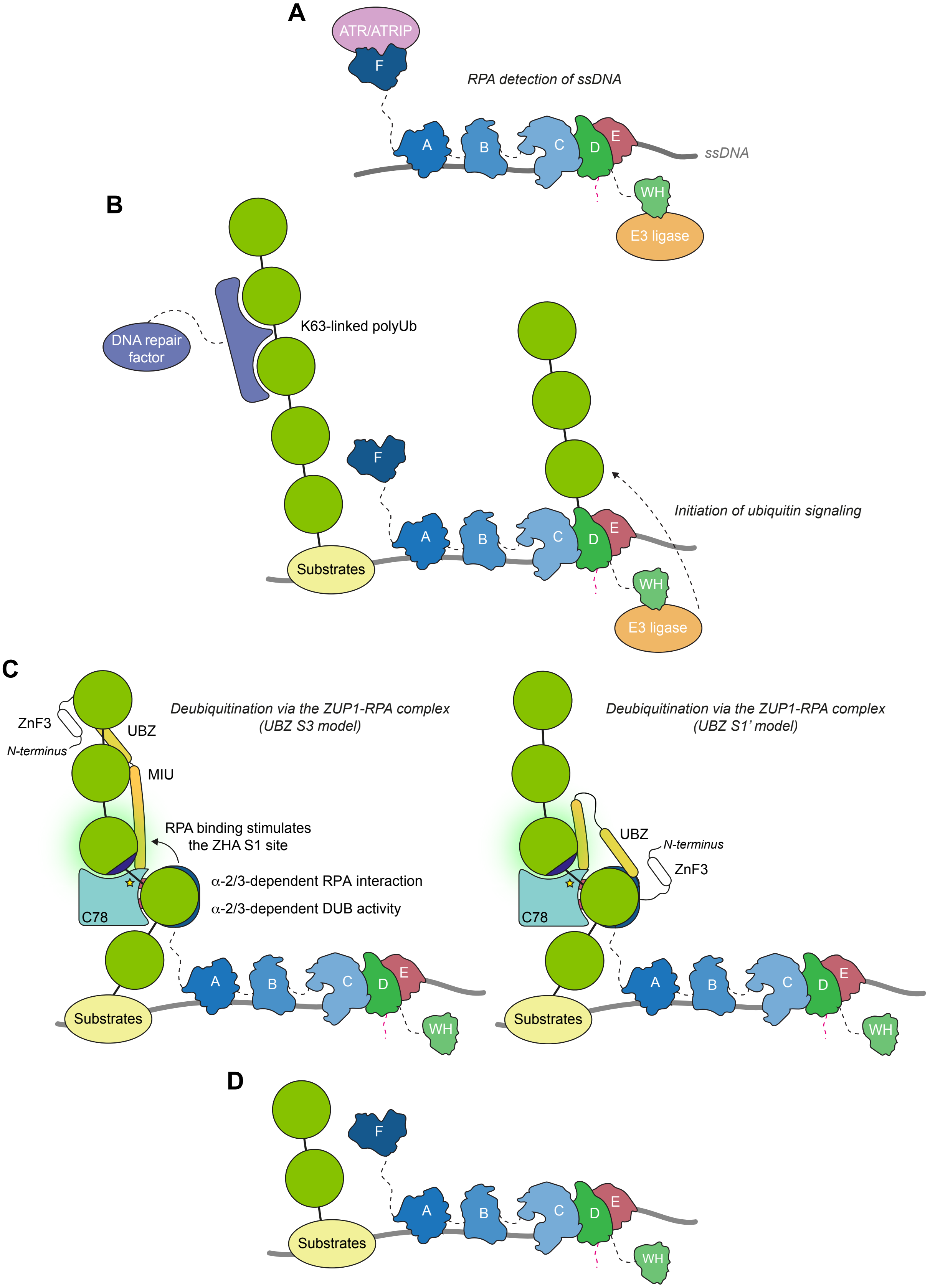

**Extended Data Figure 7. A model for ZUP1-RPA complex function.**

- **A.** ssDNA formed during DNA replication or DNA repair leads to RPA loading and the recruitment of replication and repair factors, including the ATR/ATRIP kinase complex and various E3 ligases.
- **B.** Ubiquitination of proteins, including RPA, occurs in the local ssDNA environment, leading to Ub-dependent DNA repair factor recruitment, via UBDs, and regulation.
- **C.** The ZUP1-RPA complex targets K63-linkages in long homotypic or heterotypic polyUb chains to limit their signaling capacity at ssDNA regions. Beyond RPA, the cellular substrates of the ZUP1-RPA complex are unknown. *Left*, the UBZ S3 model. *Right*, the UBZ S1’ model.
- **D.** Given the ZUP1 preference for long polyUb chains, further removal of Ub might occur by another DUB with a preference for short chains or the remaining Ub might be modified again by E3 ligases.

### Extended Data Figure S8

## A

| Supergroup | Members | Homolog? | Number | Domain arrangement | Representative |
|---|---|---|---|---|---|
| TSAR | Telonemia | No | | | |
| | Stramenopila | Yes | ≤20 | C78 or ZnF_C78 or 6xMIU_C78 | *Chaetoceros tenuissimus* |
| | Alveolata | Yes | 1 | 5xMIU_C78 | *Vitrella brassicaformis* |
| | Rhizaria | Yes | 1 | 3xZnF_C78 | *Plasmodiophora brassicae* |
| Haptista | Centrohelida | No | | | |
| | Haptophyta | No | | | |
| Cryptista | Palpitomonas | No | | | |
| | Katablepharida | No | | | |
| | Cryptophta | Yes | ≤10 | C78 | *Guillardia theta* |
| Archaeplastida | Rhodophyta | Yes | ≤10 | ZnF_MIU_C78 | *Chondrus crispus* |
| | Chloroplastida | | | | |
| | Chlorophyta | Yes | ≤25 | ZnF_C78 or MIU_C78 or 2xZnF_C78 | *Coccomyxa* |
| | Streptophyta | | | | |
| | Klebsormidiophyceae | Yes | ≤10 | C78 or 2xZnF_C78 | *Klebsormidium nitens* |
| | Phragmoplastophyta | | | | |
| | Charophyceae | No | | | |
| | Coleochaetophyceae | No | | | |
| | Zygnematophyceae | No | | | |
| | Embryophyta | Yes | ≤650 | ZnF_MIU_C78 | *Arabidopsis thaliana* |
| | Glaucophyta | No | | | |
| Ancoracysta | | No | | | |
| Picozoa | | No | | | |
| Amorphea | Apusomonada | No | | | |
| | Breviates | No | | | |
| | Opisthokonta | Yes | ≥1000 | 4xZnF_MIU_ZHA_$\alpha$2/3_C78 | *Homo sapiens* |
| | Amoebozoa | Yes | ≤10 | ZnF-C78 | *Acanthamoeba castellanii* |
| CRUMS | Diphylleida | No | | | |
| | Rigifilida | No | | | |
| | Mantamonas | No | | | |
| Excavates | Discoba | No | | | |
| | Metamonada | Yes | ≤10 | C78 | *Monocercomonoides exilis* |
| | Malawimonadida | No | | | |
| Ancyromonadida | | No | | | |
| Hemimastiophora | | No | | | |

### Extended Data Figure S8

**B**

Amorphea > Opisthokonta > Holomycota

| Group | Members | Homolog? | Number | Domain arrangement | Representative |
|---|---|---|---|---|---|
| Cristidiscoidea | | No | | | |
| Fungi | Opisthosporidia | No | | | |
| | Chytridiomycota | Yes | ≤10 | C78 or ZnF_C78 or 2xZnF_C78 | *Batrachochytrium salamandrivorans* |
| | Neocallimastigomycota | No | | | |
| | Blastocladiomycota | No | | | |
| | Zoopagomycota | Yes | ≤10 | C78 or ZnF_C78 | *Basidiobolus meristosporus* |
| | Mucoromycota | Yes | ≤150 | C78 or ZnF_C78 or 2xZnF_C78 | *Dissophora ornata* |
| | Glomeromycota | No | | | |
| | Ascomycota | | | | |
| | Pezizomycotina | | | | |
| | Laboulbeniomycetes | No | | | |
| | Arthoniomycetes | No | | | |
| | Lecanoromycetes | Yes | ≤10 | C78 or 2xZnF_C78 | *Lasallia pustulata* |
| | Leotiomyceta | Yes | ≤140 | ZnF_C78 | *Diplocarpon rosae* |
| | Dothideomyceta | Yes | ≤360 | 2xZnF_C78 | *Aureobasidium melanogenum* |
| | Sordariomyceta | Yes | ≤550 | C78 or 2xZnF_C78 | *Colletotrichum higginsianum* |
| | Pezizomycetes | Yes | ≤20 | 2xZnF_C78 | *Tuber magnatum* |
| | Orbiliomycetes | Yes | ≤20 | C78 or 2xZnF_C78 | *Orbilia oligospora* |
| | Eurotiomycetes | Yes | ≤250 | 2xZnF_C78 | *Aspergillus niger* |
| | Saccharomycotina | No | | | |
| | Taphrinomycotina | Yes | ≤10 | C78 or ZnF_C78 or 2xZnF_C78 | *Schizosaccharomyces pombe* |
| | Basidiomycota | | | | |
| | Pucciniomycotina | Yes | ≤50 | 2xZnF_C78 | *Atractiella rhizophila* |
| | Orthomycotina | | | | |
| | Ustilaginomycotina | No | | | |
| | Agaricomycotina | | | | |
| | Wallemiomycetes | No | | | |
| | Tremellomycetes | Yes | ≤70 | 2xZnF_C78 | *Kockovaella imperatae* |
| | Dacrymycetes | Yes | ≤10 | ZnF_C78 | *Calocera cornea* |
| | Agaricomycetes | Yes | ≤320 | C78 or ZnF_C78 | *Auriscalpium vulgare* |

**Extended Data Figure S8**

**C**

Amorphea > Opisthokonta > Holozoa

| Group | Members | Homology? | Number | Domain arrangement | Representative |
|---|---|---|---|---|---|
| Mesomycetozoea | | No | | | |
| Corallochytrium | | No | | | |
| Filozoa | Filasterea | No | | | |
| | Choanoflagellata | Yes | ≤5 | ZnF_C78 | *Monosiga brevicollis* |
| | Animalia | | | | |
| | Porifera | No | | | |
| | Eumetazoa | | | | |
| | Ctenophora | No | | | |
| | Parahoxozoa | | | | |
| | Cnidaria | Yes | ≤20 | 2xZnF_MIU_ZHA_α2/3_C78 | *Nematostella vectensis* |
| | Placozoa | Yes | ≤5 | 2xZnF_α2/3_C78 | *Trichoplax adhaerens* |
| | Bilateria | | | | |
| | Xenacoelomorpha | No | | | |
| | Nephrozoa | | | | |
| | Deuterostomia | | | | |
| | Chordata | | | | |
| | Cephalochordate | Yes | ≤5 | 4xZnF_MIU_ZHA_α2/3_C78 | *Branchiostoma floridae* |
| | Olfactores | | | | |
| | Tunicata | Yes | ≤5 | 3xZnF_MIU_ZHA_α2/3_C78 | *Styela clava* |
| | Vertebrata | | | | |
| | Fish | Yes | ≤350 | 4xZnF_MIU_ZHA_α2/3_C78 | *Danio rerio* |
| | Amphibians | Yes | ≤20 | 4xZnF_MIU_ZHA_α2/3_C78 | *Xenopus laevis* |
| | Reptiles | Yes | ≤30 | 4xZnF_MIU_ZHA_α2/3_C78 | *Chelydra serpentina* |
| | Birds | Yes | ≤200 | 4xZnF_MIU_ZHA_α2/3_C78 | *Gallus gallus* |
| | Mammals | Yes | ≤200 | 4xZnF_MIU_ZHA_α2/3_C78 | *Homo sapiens* |
| | Ambulacraria | | | | |
| | Echinodermata | Yes | ≤10 | 4xZnF_MIU_ZHA_α2/3_C78 | *Strongylocentrotus purpuratus* |
| | Hemichordata | No | | | |
| | Protostomia | | | | |
| | Ecdysozoa | | | | |
| | Scalidophora | Yes | ≤5 | 4xZnF_ZHA_α2/3_C78 | *Priapulus caudatus* |
| | Panarthropoda | | | | |
| | Onychophora | No | | | |
| | Tardigrada | Yes | ≤5 | ZnF_2xMIU_C78 | *Hypsibius exemplaris* |
| | Arthropoda | Yes | | | |
| | Chelicerates | Yes | ≤25 | 5xZnF_MIU_ZHA_α2/3_C78 | *Centruroides sculpturatus* |
| | Myriapods | No | | | |
| | Pancrustaceans | | | | |
| | Oligostraca | Yes | ≤5 | 3xZnF_MIU_ZHA_α2/3_C78 | *Darwinula stevensoni* |
| | Multicrustacea | Yes | ≤25 | 5xZnF_ZHA_α2/3_C78 | *Portunus trituberculatus* |
| | Allotriocarida | | | | |
| | Branchiopoda | Yes | ≤10 | 3xZnF_MIU_ZHA_α2/3_C78 | *Daphnia magna* |
| | Cephalocarida | No | | | |
| | Remipedia | No | | | |
| | Hexapoda | Yes | ≤370 | 5xZnF_ZHA_α2/3_C78 | *Tribolium castaneum* |
| | Nematoida | Yes | ≤5 | 3xZnF_C78 | *Toxocara canis* |
| | Spiralia | | | | |
| | Gnathifera | Yes | ≤30 | 2xZnF_MIU_ZHA_α2/3_C78 | *Adineta vaga* |
| | Lophotrochozoa | | | | |
| | Platyhelminthes | Yes | ≤45 | ZnF_MIU_ZHA_α2/3_C78 | *Macrostomum lignano* |
| | Nemertea | No | | | |
| | Annelida | Yes | ≤5 | 5xZnF_ZHA_α2/3_C78 | *Owenia fusiformis* |
| | Phoronida | No | | | |
| | Ectoprocta | No | | | |
| | Brachiopoda | Yes | ≤5 | 5xZnF_ZHA_α2/3_C78 | *Lingula anatina* |
| | Gastrotricha | No | | | |
| | Mollusca | Yes | ≤30 | 5xZnF_ZHA_α2/3_C78 | *Lottia gigantea* |
| | Entoprocta | No | | | |

**Extended Data Figure 8. Distribution of ZUP1 homologs across eukaryotes.**

- **A.** A catalogue of ZUP1 homologs in the eukaryotic supergroups, with the indicated number of species, the domain arrangement, and a representative member. Homologs were identified using BLAST with *Hs*ZUP1 as a query. Domains were identified via a combination of analysing the BLAST alignments, by manual inspection, and from AlphaFold predictions.
- **B.** As in (A), except more detail for Amorphea > Opisthokonta > Holomycota.
- **C.** As in (A), except more detail for Amorphea > Opisthokonta > Holozoa.
